# Evolution and Ecology in Widespread Acoustic Signaling Behavior Across Fishes

**DOI:** 10.1101/2020.09.14.296335

**Authors:** Aaron N. Rice, Stacy C. Farina, Andrea J. Makowski, Ingrid M. Kaatz, Philip S. Lobel, William E. Bemis, Andrew H. Bass

## Abstract

Acoustic signaling by fishes has been recognized for millennia, but is typically regarded as comparatively rare within ray-finned fishes; as such, it has yet to be integrated into broader concepts of vertebrate evolution. We map the most comprehensive data set of volitional sound production of ray-finned fishes (Actinopterygii) yet assembled onto a family level phylogeny of the group, a clade representing more than half of extant vertebrate species. Our choice of family-level rather than species-level analysis allows broad investigation of sonifery within actinopterygians and provides a conservative estimate of the distribution and ancestry of a character that is likely far more widespread than currently known. The results show that families with members exhibiting soniferous behavior contain nearly two-thirds of actinopterygian species, with potentially more than 20,000 species using acoustic communication. Sonic fish families also contain more extant species than those without sounds. Evolutionary analysis shows that sound production is an ancient behavior because it is present in a clade that originating circa 340 Ma, much earlier than any evidence for sound production within tetrapods. Ancestral state reconstruction indicates that sound production is not ancestral for actinopterygians; instead, it independently evolved at least 27 times, compared to six within tetrapods. This likely represents an underestimate for actinopterygians that will change as sonifery is recognized in ever more species of actinopterygians. Several important ecological factors are significantly correlated with sonifery – including physical attributes of the environment, predation by members of other vertebrate clades, and reproductive tactics – further demonstrating the broader importance of sound production in the life history evolution of fishes. These findings offer a new perspective on the role of sound production and acousticcommunication during the evolution of Actinopterygii, a clade containing more than 34,000 species of extant vertebrates.

## Introduction

While spoken language is regarded as a uniquely human attribute, the use of sound as a vertebrate communication channel also occurs in other terrestrial species and marine mammals (Bradbury and Vehrencamp 2011, Ladich and Winkler 2017). Less well known is its prevalence among fishes, despite multiple early descriptions of anatomy, physiology or behavior (Dufossé 1874, Tower 1908), including von Frisch’s comments on its widespread distribution as early as 1938:

> It may well be asked for what purpose fishes are able to hear so well in silent water. … We know many species of sound-producing fish. There may be many more species of sound-producing fishes not yet known. … [and] much to discover in the future about the language of fishes. (von Frisch 1938)

Since then, a growing body of evidence shows the importance of volitional sound production in social communication and reproduction especially among ray-finned fishes (Actinopterygii) (Ladich 2015), a group that includes more than half of extant vertebrate diversity. Together with Sarcopterygii (coelacanths, lungfishes, and tetrapods, which includes amphibians, reptiles, birds, and mammals), Actinopterygii is one of two extant radiations of bony vertebrates(Nelson et al. 2016). Although there is evidence for soniferous behavior in 800-1000 species of actinopterygians (Ladich 2015, Ladich et al. 2006) and numerous studies of neural and hormonal mechanisms that are similar to those of tetrapods (Bass 2014, Zhang and Ghazanfar 2020), more widespread recognition of acoustic behavior among fishes and its integration into broader concepts of vertebrate evolution are still lacking. This is, in part, because sound production is not externally obvious in fishes, nor can those sounds be easily detected underwater without specialized technology (Mann et al. 2016).

A recent study on the evolution of acoustic communication focused on tetrapods, recognized the important need for a comparable study of fishes (Chen and Wiens 2020). Using evolutionary modelling, combined with the most recent comprehensive phylogeny, we show that volitional sound production is ancestral for several speciose radiations that together comprise nearly two-thirds of the 34,000 valid extant species of actinopterygians (Fricke et al. 2020). We also show that sound production has evolved at least 27 times among actinopterygians, including the basal clade that diverged in the Carboniferous Period (∼340 Ma). Thus, actinopterygian sonifery is likely an ancient communication mode that originated earlier than estimates for the origin of acoustic communication in tetrapods where it is proposed to have evolved six times (Chen and Wiens 2020). Nocturnality was identified as the one ecological factor contributing to the evolution of acoustic communication among tetrapods (Chen and Wiens 2020). We show that actinopterygian families with soniferous species are correlated with multiple ecological factors, including reproductive and mating tactics, trophic levels and complexity of habitats that vary in depth, substrate composition, and salinity.

In aggregate, our evidence strongly supports the hypothesis that, like tetrapods, acoustic communication is an ancient but also convergently evolved innovation across actinopterygian fishes. Unlike tetrapods, we find that actinopterygian soniferous behavior is associated with a broad range of abiotic and biotic factors, which may explain its repeated and independent evolution nearly 30 times in clades that include many of the most species-rich groups. The demonstration of repeated evolution of acoustic communication in tetrapods and now in ray-finned fishes highlights the strong selection pressure favoring this signaling modality across vertebrates.

## Materials and Methods

We operationally define acoustic signaling, or soniferous behavior (we use these terms interchangeably) as volitional sound production associated with acoustic communication rather than by-products of feeding or locomotion. Like Chen and Wiens (2020), we score the presence or absence of soniferous behavior at a family level, in this case for valid extant species of actinopterygians in 461 families represented by species in Rabosky et al. (2018) with the assumption that sonifery is conserved and characteristic at the family level (Fricke et al. 2018). We use three lines of evidence from one or more reports to demonstrate the presence of soniferous behavior in 167 of the 461 families in our analysis (Fig. S1, Tables S1, S2): 1) quantitative or pictorial documentation of acoustic recordings (107 families); 2) the presence of specialized morphology strongly predictive of sonic ability (Fine and Parmentier 2015) (26 families); or 3) qualitative descriptions of sounds strongly predictive of sonic ability and behaviorally-relevant acoustic signals (Hubbs 1920, von Frisch 1938) (34 families). To be conservative, we code as 0 (silent) all families lacking such evidence.

**Table 1.** Probabilities sound production is ancestral state, and phylogenetic signal for Actinopterygii (ray-finned fishes) and some of its sub-clades

| Clade | Number of Extant Families in Figure 2 | Estimated Number of Valid Extant Species<sup>1</sup> | Probability Soniferous Behavior is Ancestral to Clade<sup>2,3</sup> | Phylogenetic Signal |  |
| --- | --- | --- | --- | --- | --- |
|  |  |  |  | D statistic | Probability of Brownian Phylogenetic Structure<sup>2</sup> |
| Actinopterygii | 461 | 34,030 | 29.4% | 0.404 | 0.037 |
| Teleostei | 456 | 33,970 | 15.2% | 0.368 | 0.057 |
| Osteoglossomorpha | 6 | 250 | 25.5% | 1.680 | 0.165 |
| Otocephala | 96 | 11,720 | 9.6% | 0.328 | 0.288 |
| Ostariophysi | 88 | 11,160 | 8.5% | 0.208 | 0.400 |
| Curimatoidea | 6 | 420 | 67.0% | -31.388 | 0.809 |
| Siluroidei | 30 | 2,340 | 96.7% | -0.469 | 0.609 |
| Euteleostei | 333 | 20,930 | 9.6% | 0.200 | 0.231 |
| Acanthomorpha | 298 | 19,470 | 31.4% | 0.270 | 0.164 |
| Eupercaria | 142 | 6,970 | 88.6% | -0.066 | 0.589 |
| Anabantaria + Carangaria + Ovalentaria | 81 | 7,300 | 64.1% | 0.075 | 0.489 |
| Hexagrammidae + Zoarcoidei + Cottoidei | 25 | 1,280 | 97.9% | -0.676 | 0.760 |
| Cottoidei | 11 | 850 | 98.8% | -0.330 | 0.644 |

<sup>1</sup> Rounded to nearest 10 based on 04 April 2020 download of (Fricke et al. 2020).
<sup>2</sup> Probabilities for ancestral state and Brownian phylogenetic structure are represented differently to help distinguish them.
<sup>3</sup> Node percentages summarize 1000 stochastic character mapping simulations.

Data on fish sound production were obtained from journals, technical reports, conference proceedings, theses, and books (Table S1). We mapped the presence (= 1) or absence (i.e. silent, = 0) of soniferous behavior onto Rabosky et al.’s (2018) recent phylogeny of Actinopterygii that includes species from 461 families (Fig. S1, Table S1). Species included in the phylogeny by Rabosky et al. (2018) were assigned to families using Catalog of Fishes (Fricke et al. 2018).

Since Rabosky et al. (2018), new species have been described and familial designations changed (Fricke et al. 2020). We note that four families in our analyses (Abyssocottidae, Comephoridae, Cynolebiidae, Hapalogenyidae) were merged into other families, and approximately 14 new families were recognized (Fricke et al. 2020).

We scored the presence or absence of soniferous behavior as a binary character (Table S2). Ancestral states were calculated using stochastic character mapping with the make.simmap function in the *phytools* (Revell 2012) package for R, with 1,000 MCMC generations, sampling every 100 generations. Root node values and transition rates were calculated by simulation and posterior probabilities were mapped using the densityMap function in *phytools* (Revell 2012) (Figs 1, 2). Phylogenetic signal was calculated using the *D* statistic (Fritz and Purvis 2010) with the *caper* R package (Orme et al. 2013).

Ecological attributes for all 461 families were downloaded from FishBase(Froese and Pauly 2019) using *rfishbase* 3.04 R package (Boettiger et al. 2012) (see SI Appendix, Table S2 for complete data). Ecological parameters predictive of soniferous behavior were determined using logistic regression with a phylogenetic generalized linear model (Ives and Garland 2010) in *phylolm* 2.6 R package (Ho and Ané 2014). Since we tested several models for each set of parameters, we used Bonferroni correction to reduce Type I error (Rice 1989). Data on species number per family are from the Eschmeyer Catalog of Fishes (Fricke et al. 2020).

## Results

### Ancestral States

Stochastic character mapping simulates the distribution of a character along branches of a phylogeny (Bollback 2006, Revell 2012) and summaries of many simulations (N = 1000 in this study) are used to compute probabilities of a character being ancestral at nodes.

Figure 1 reconstructs ancestral states of soniferous behavior across actinopterygian phylogeny, showing the probabilities of soniferous behavior being ancestral, ranging from 0% (silent) to 100% (soniferous); Table 1 presents probability values at key nodes.

**Fig. 1.**
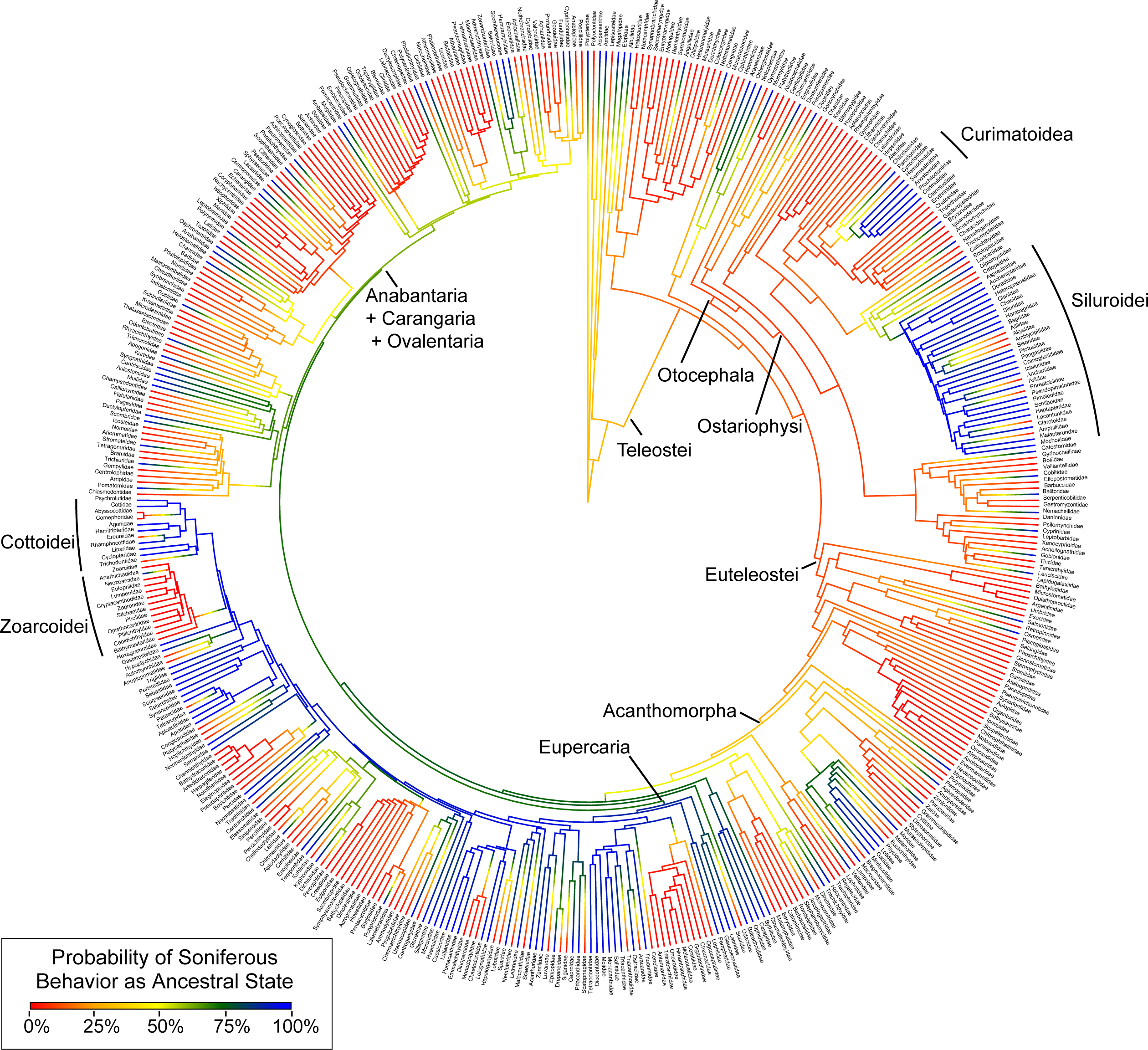
Family-level phylogenetic tree of actinopterygians depicting evolution of soniferous behavior. Shown here are posterior probabilities from ancestral state reconstruction using stochastic character mapping. Probability is represented as a gradient, where blue indicates a high probability and red a low probability of soniferous behavior, and yellow is equivocal. Tree is pruned from species-level phylogeny of Rabosky et al.(Rabosky et al. 2018) to family-level here.

Although sonifery occurs in the three extant clades of non-teleostean actinopterygians (Polypteriformes, Acipenseriformes, and Holostei) (Fig. 1), this reconstruction reveals that soniferous behavior is unlikely ancestral for Actinopterygii (29.4% probability). Teleostei, which comprises > 99.8% of actinopterygian species, also has low support (15.2% probability) that soniferous behavior is the ancestral state. Likewise, Osteoglossomorpha, an early diverging clade of teleosts, contains several soniferous families, but only a 25.5% probability that soniferous behavior is ancestral. Otocephala, a speciose subclade of actinopterygians exhibiting morphological adaptations to enhance hearing (Braun and Grande 2008), has an even lower probability that soniferous behavior is ancestral, 9.6%. Ostariophysi, a large subgroup of otocephalans well known for the Weberian apparatus (chain of bony elements that enhance hearing), has the lowest value among the groups analyzed that soniferous behavior is ancestral, 8.5%. A second large subclade of Teleostei, Euteleostei, includes two-thirds of living fish species, but here, too, there is little support that soniferous behavior is ancestral, 9.6%.

We find much stronger support for soniferous behavior as a character at the base of some key nodes. Siluroidei, a subclade of catfishes, and Curimatoidea, a subclade of characins, have 96.7% and 67% probabilities, respectively, that soniferous behavior is ancestral (Figs. 1, 2a.

**Fig. 2.**
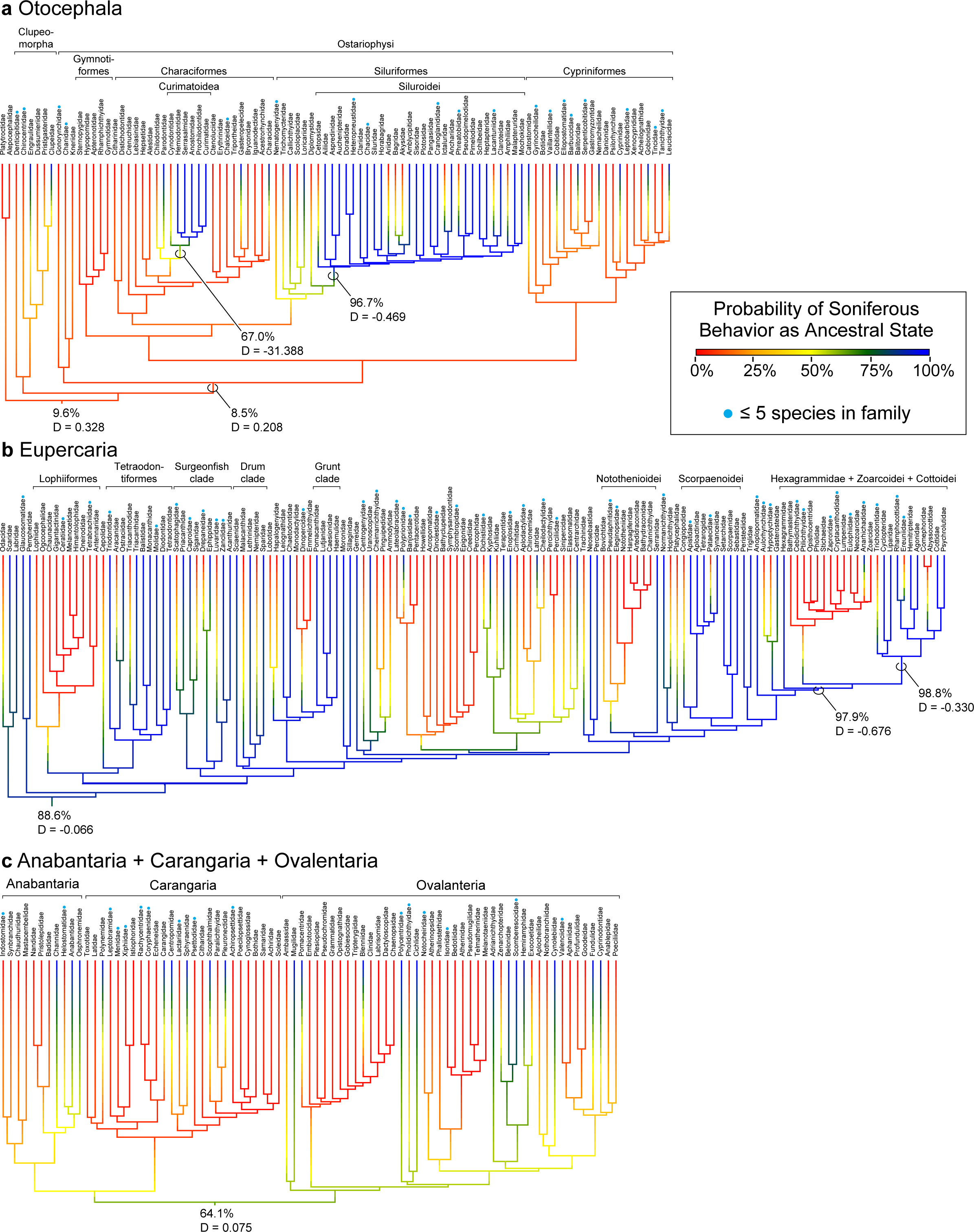
Posterior probability soniferous behavior within major actinopterygian clades. (**a**) Otocephala, (**b**) Anabantaria + Carangaria + Ovalentaria, and (**c**) Eupercaria. For phylogenetic trees showing the ancestral state estimation and associated evolutionary probabilities of sound production being ancestral by stochastic character mapping, probability is represented as a gradient where blue indicates high and red is low probability of sound production; yellow is equivocal.

Acanthomorpha, which includes 85% of fish species in marine habitats (Wainwright and Longo 2017), has a low probability (31.4%) sonifery is ancestral. However, two of its subclades, Eupercaria (e.g., “surgeonfish”, “drums”, “grunts”, scorpaenoids) and Anabantaria + Carangaria + Ovalentaria (e.g., gouramis [Osphronemidae], jacks [Carangidae], cichlids [Cichlidae]) have 88.6% and 64.1% probabilities, respectively (Fig. 2b, c). An even higher probability value, 97.5%, supports soniferous behavior as ancestral for a crown group within Eupercaria, Hexagrammidae (greenlings) + Zoarcoidei (e.g., wolffishes) + Cottoidei (e.g., sculpin) (Fig. 2b).

In aggregate, our results indicate that acoustic signaling, or soniferous behavior, has a high probability (>75%) of being ancestral for at least 27 nodes across Actinopterygii (Fig. S2). We interpret this as evidence of widespread, independent evolution of volitional sound production.

### Phylogenetic signal

Patterns of ancestral states alone do not predict evolutionary processes underlying character evolution, making it necessary to evaluate phylogenetic signal (Blomberg et al. 2003). We use the *D* statistic for binary characters (Chen and Wiens 2020, Fritz and Purvis 2010), in this case soniferous or silent, to calculate phylogenetic signal. For each clade, we computed *D* and the probability that character evolution results from Brownian phylogenetic structure, which can be visualized by the proximity of the clade’s observed *D*-value to the center of the distribution of simulated *D*-values assuming Brownian evolutionary processes (Fig. S3).

Where *D* is > 0.0, the evolution of soniferous behavior is phylogenetically random and not conserved within a group. Where *D* is close to or < 0.0, evolution of soniferous behavior results primarily from Brownian evolutionary processes and phylogenetic structure, and is conserved within a group.

Actinopterygii and Teleostei have *D* values of 0.404 and 0.368, respectively (see Table 1 for all *D* values). The next set of large clades, Otocephala, Ostariophysi and Euteleostei, have *D* values of 0.328, 0.208, and 0.200, respectively. These values indicate that soniferous behavior is not conserved within these groups, in agreement with the relatively low to intermediate probabilities that it is ancestral for these groups (8.5% - 29.4%; Table 1). For Siluroidei, a large subclade of Otocephala, *D* is -0.469, consistent with the high probability that this character is ancestral for Otocephala (96.7%, Table 1).

Acanthomorpha has *D* = 0.270, in agreement with the relatively low probability that sonifery is ancestral for this group (Table 1). However, within Acanthomorpha, several nested groups show negative *D* values or values very close to 0.0, in agreement with the high probabilities that soniferous behavior is ancestral for these groups (Table 1). This includes two large acanthomorph clades, Eupercaria and Anabantaria + Carangaria + Ovalentaria, with *D* values of -0.066 and 0.075, respectively. Within Eupercaria, Hexagrammidae + Zoarcoidei + Cottoidei, *D* = -0.676. The two smallest subclades studied, Osteoglossomorpha and Curimatoidea, have *D* values of 1.680 and -31.388, respectively, that agree with low (Osteoglossomorpha) and high (Curimatoidea) probabilities sonifery is ancestral for these groups (Table 1).

### Hearing specializations

Novel auditory morphologies, generally referred to as hearing specializations, e.g., the Weberian apparatus or swim bladder extensions contacting the otic capsule, may have evolved 20 times within Teleostei (Braun and Grande 2008). Families with these adaptations (Braun and Grande 2008, Colleye et al. 2019, Radford et al. 2013) (Table S2), 62 of 119, are highly correlated with soniferous behavior (phylogenetic logistic regression; *P* = 0.004).

### Habitat Complexity

Actinopterygian families with soniferous taxa live in habitats that vary in complexity depending on one or more of the following: water salinity, depth and substrate composition (Boettiger et al. 2012, Froese and Pauly 2019) (Table S2). Freshwater and brackish water are more likely than marine habitats to have families with soniferous taxa (*P* < 0.000, < 0.000, > 0.05, respectively; values here and below based on logistic regression with a phylogenetic generalized linear model(Ives and Garland 2010) after Bonferroni correction). Marine families in shallow intertidal (< 5 m depth) and neritic (< 200 m depth) zones are more likely to have soniferous taxa (*P* < 0.000) than families with oceanic (i.e. marine pelagic) fishes (*P* > 0.05). Within families with freshwater species, there is no significant correlation of soniferous behavior with depth (littoral zone, sublittoral zone, caves; *P* values > 0.05). Habitats with coarse (*P* = 0.008), but not fine (*P* > 0.05), sediment are also more likely to have families with soniferous taxa. Soniferous families are not more likely to live in any one particular climate (polar, temperate, boreal, tropical, subtropical; *P* values > 0.05).

Grosberg et al. (2012) consider the complexity of freshwater and marine environments, and how more structurally complex habitats are associated with higher biodiversity. Of the 27 independent evolutionary events of soniferous behavior we describe (Fig. S2, Table S3), 18 clades are primarily freshwater, and nine are either marine, anadromous, or mixed. With the exception of Myctophidae, 26 of the 27 clades live in shallow waters or demersal/benthic habitats.

### Feeding and Reproductive Ecologies

Actinopterygian families exhibiting acoustic signaling are associated with several other behavioral phenotypes (Table S2). Marine families with grazing species are more likely to contain soniferous taxa (*P* = 0.011), as are families with mating tactics and reproductive modes ranging from batch spawning (*P* < 0.0001) and internal fertilization (*P* = 0.005), to nest guarding (*P* = 0.001), parental care (*P* = 0.004) and alternative reproductive tactics (17 of 23 families identified by Mank and Avise (2006); *P* < 0.0001). Families showing sex reversal (protogyny, protandry, hermaphroditism) are not more likely to contain soniferous taxa (P > 0.05). Consistent with field observations, actinopterygian families with soniferous taxa are significant prey for birds (Elliott et al. 2003) and elasmobranchs (Navia et al. 2007) (*P* = 0.002, 0.001, respectively; cetaceans (McCabe et al. 2010) and pinnipeds(Lance and Jeffries 2009) are known predators, but *P* values > 0.05).

## Discussion

Although actinopterygian fishes have long been known capable of volitional sound production (Popper and Casper 2011), few studies integrate their acoustic communication ability into a broad evolutionary context across bony vertebrates (Bass et al. 2015, Fine and Parmentier 2015). We show evidence for soniferous behavior in 167 families, containing nearly two-thirds of the estimated 34,000 valid extant species of actinopterygians (Figs. 1, 2; Tables S1, S2).

Actinopterygians independently evolved soniferous ability at least 27 times (Fig. S2, Table S3). To our knowledge, all species studied to date that are capable of volitional sound production have been shown to use sound in a signaling context to either conspecific or heterospecific individuals (Ladich 2015, Ladich et al. 2006). Consequently, sound production is likely an important communication modality in most actinopterygian species. This includes two species of polypterids (Ladich and Tadler 1988), members of a family that diverged from the actinopterygian stem circa 340 Ma during the Carboniferous Period (Giles et al. 2017). This suggests that acoustic communication in actinopterygians may have similarly ancient origins, predating its emergence within tetrapods, which occurred circa 100-200 Ma (Chen and Wiens 2020). We further show significant correlations between families with soniferous species and diverse freshwater and marine habitats, predation by birds and elasmobranchs, and many reproductive and mating tactics. In parallel with recent findings for tetrapods(Chen and Wiens 2020), our results indicate strong selection to exploit acoustic signaling for communication and ecological success across vertebrate evolution.

### Pattern and process

Within Actinopterygii, soniferous behavior occurs across the most speciose clades and has evolved independently at least 27 times, compared to only six within tetrapods (Chen and Wiens 2020). This high frequency of convergent evolution suggests that “the interplay of historical contingency and natural selection” (Blount et al. 2018) has a prominent role in the evolution of vertebrate acoustic communication behavior. A comparable degree of convergent evolution among actinopterygians is reported for alternative reproductive tactics (Mank and Avise 2006), suggesting that extensive convergence may be an evolutionary attribute of behavioral and reproductive ecology as well as other characters in actinopterygians (e.g., venom (Smith and Wheeler 2006), restricted gill openings (Farina et al. 2015), vertebrae (Ward and Brainerd 2007), adipose fins (Stewart et al. 2014), migratory behavior (Burns and Bloom 2020), bioluminescence (Davis et al. 2014)).

The presence and absence of soniferous behavior among actinopterygians likely includes secondary loss, suggested elsewhere to drive speciation (Miles and Fuxjager 2019). Within speciose clades where sonifery has a high probability of being ancestral (Siluroidei, Eupercaria, Anabantaria + Carangaria + Ovalentaria, Hexagrammidae + Zoarcoidei + Cottoidei), non-soniferous clades may have secondarily lost this character. Hexagrammidae + Zoarcoidei + Cottoidei have 97.9% probability that sound production is ancestral, and a very low *D* value (- 0.676, Table 1, Fig. 2b). Within this group, Cottoidei comprises an estimated 850 species (compared to 9 hexagrammid and 405 zoarcoid species) with a very high probability that soniferous behavior is ancestral (98.8%). This correlates with a low *D* value (-0.330), suggesting that the evolution of soniferous behavior within Cottoidei results primarily from Brownian evolutionary processes. Fish and Mowbray (1970) comment on the absence of sound production in Zoarcidae [their Zoarchidae]. If further research provides conclusive evidence for absence, then our tree (Fig. 2c) likely indicates secondary loss. Other places to investigate potential loss of soniferous capacity are between sister groups where one is coded as silent (e.g., Lophiiformes) and the other is soniferous (Tetraodontiformes; Fig. 2c). A particularly fascinating case of secondary loss concerns catfishes in the genus *Synodontis*; some species are only soniferous and others only weakly electric (Boyle et al. 2014). Weakly electric *Synodontis* have reduced sonic muscle characters, but share characters with myogenic electric organs (Kéver et al. 2020).

Further demonstration of the loss of sonifery would support the hypothesis that losses can be as important in generating diversity as gains of complexity (Miles and Fuxjager 2019).

Together, *D* values show soniferous behavior is highly conserved (low *D*) in some lineages, but less in others (high *D*). Comparisons of ancestral state probabilities and *D* values show that clades with a higher probability of soniferous ability in the common ancestor also tend to have lower *D* values (Table 1, Fig. 3a), indicating that when it is ancestral, it has a higher probability of being conserved within a clade. This relationship becomes even clearer when plotting ancestral state probabilities against the probability that phylogenetic signal results from Brownian phylogenetic structure (Fig. 3b). It may intuitively follow that an ancestral trait is more likely to be conserved, but these two metrics are independently derived.

**Fig. 3.**
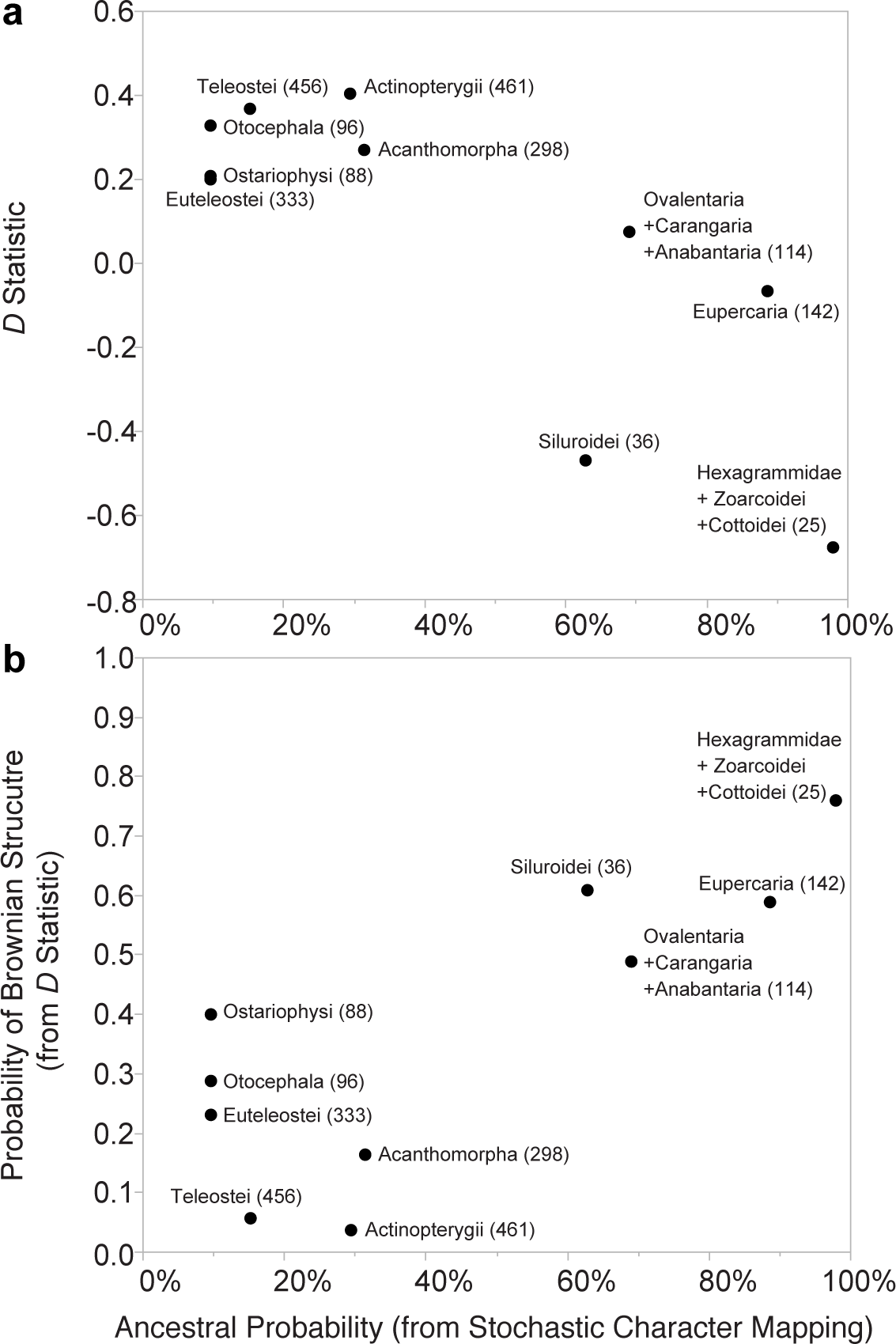
Phylogenetic signal versus ancestral state estimation for evolution of actinopterygian soniferous behavior. (**a**) *D* statistic value (Fritz and Purvis 2010) versus ancestral state estimate (using stochastic character mapping) probability that soniferous behavior is ancestral for a clade. (**b**) Probability of Brownian phylogenetic structure (modelled from *D* statistic) versus stochastic character mapping probability soniferous behavior is ancestral for a clade. Values for *D* statistic, probability of Brownian structure, and ancestral state probabilities are listed in Table 1. Only clades with >25 families are used, since inference of *D* is limited for clades with <25 taxa (Fritz and Purvis 2010).

Plotting the relationship between ancestral state estimation and phylogenetic signal may indicate a broader conceptual link between pattern (ancestral states) and process (phylogenetic signal) in character evolution (Fig. 3b). Some cases deviate from this relationship. For example, it is unlikely that soniferous behavior is ancestral for Ostariophysi, yet the character is relatively conserved within this clade. Exceptions indicate that the relationship is not necessitated mathematically, but instead is governed by evolutionary principles. Characters that vary enormously in phylogenetic signal throughout lineages and are characterized by repeated gains and losses, such as soniferous behavior, may be more likely to exhibit a relationship between ancestral state and phylogenetic signal.

### Ecological success

Our results provide compelling evidence that soniferous evolution contributes to ecological success in many actinopterygian clades, as it does in tetrapods and insects (Miles et al. 2018, Wilkins et al. 2013). For example, we can now add soniferous behavior to the suite of traits considered as evolutionary drivers in Acanthomorpha, which account for 85% of fish species in marine habitats (Wainwright and Longo 2017), because many soniferous species belong to basal acanthomorph groups, e.g., Beryciformes, Ophidiiformes, and Gadiformes (Fig. 1). Soniferous behavior may be a convergent evolutionary innovation contributing to ecological success in rapidly evolving and speciose subclades of actinopterygians for which sonifery is ancestral. For example, Eupercaria and Siluroidei are nested within rapidly evolving lineages in Actinopterygii (Table 1) (Alfaro et al. 2009), and it is intriguing to hypothesize that sonifery may promote diversification through sexual selection. This also appears to be the case for soniferous tetrapods, including birds and eutherian mammals (Alfaro et al. 2009, Chen and Wiens 2020).

Molecular phylogenetic support for Curimatoidea, a clade recently recognized (Arcila et al. 2017, Betancur-R. et al. 2019) within Characiformes (Figs 1, 2), is bolstered by our evidence that soniferous behavior is ancestral for this clade. Intriguingly, the relationship between repeated evolution of soniferous behavior in clades that live in shallow water or structurally complex or fragmented habitats where diversification is more likely to occur (Grosberg et al. 2012), suggests a strong selection for acoustic communication within biodiverse communities.

Urick (1975) points out at the very beginning of his classic text, Principles of Underwater Sound, that water is an excellent medium for sound transmission compared to other modalities:

> Of all the forms of radiation … sound travels through the sea the best. In the turbid, saline water of the sea, both light and radio waves are attenuated to a far greater degree than is that form of mechanical energy known as sound. (Urick 1975)

The relationship between physical sound transmission in an aquatic medium with acoustic communication has previously been identified as promoting this modality in underwater habitats (Grosberg et al. 2012, Wilkins et al. 2013). Our analyses show that sonifery is correlated with families living in fresh or brackish waters, marine intertidal or neritic zones, and habitats with coarse as opposed to fine sediment bottoms. Salinity, water depth and substrate composition are all physical properties of the environment that impact acoustic properties (Forrest et al. 1993, Urick 1975). For example, transmission loss due to absorption (“conversion of acoustic energy into heat”; Urick 1975) is greater in seawater and shallow water. Sound speed is greater in bottoms with coarse substrates, but unpredictable in shallow water because of salinity, currents and changes in temperature at the surface. To more completely understand how physical properties of the environment combine to impact acoustic communication, direct measurements are needed in a range of habitats (Bass and Clark 2003, Lugli 2015).

We report a correlation between families exhibiting soniferous behavior and hearing specializations that enhance sound detection. This enhances the efficacy of other physiological mechanisms for audio-vocal coupling that support acoustic communication. Actinopterygians share with tetrapods (and insects) two hallmarks of audio-vocal coupling: auditory encoding of the spectral and temporal properties of conspecific and heterospecific vocalizations (Bass et al. 2005, Rohmann et al. 2013), and a central vocal corollary discharge, whereby vocal pattern generator neurons inform auditory neurons about the spectral and temporal attributes of one’s own vocalizations (Chagnaud and Bass 2013).

Perhaps the most compelling evidence that acoustic signaling behavior contributes to ecological success within Actinopterygii is the evidence we present of its association with alternative mating tactics and multiple modes of reproduction, including nest guarding, batch spawning, internal fertilization, and parental care. These findings point to many taxa of soniferous actinopterygians as providing new testing grounds for investigating the influence of sexual and ecological selection, and drift on the evolution of acoustic communication systems (Amorim et al. 2018, Bose et al. 2018, Emlen and Oring 1977, Lee and Bass 2006, Myrberg and Riggio 1985, Wilkins et al. 2013).

### Concluding Comments

The remarkable ecological, behavioral, and morphological diversity of actinopterygian fishes provides opportunities to test evolutionary trajectories, constraints or roles of acoustic communication. Because several key functional innovations have been associated with diversification and evolutionary success in actinopterygians (e.g., acanthomorphs; Wainwright and Longo 2017), we argue that sound production and acoustic signaling may be similar key innovations in actinopterygian evolution. In a broader sense, and together with recent demonstrations of acoustic communication in tetrapods (Chen and Wiens 2020), our findings highlight the important role that acoustic communication has played in the history of vertebrates.

## Acknowledgements

Research supported, in part, by National Science Foundation awards OCE-1736936 (ANR), DBI-1523836 (SCF), and IOS-1656664 (AHB), the Tontogany Creek Fund (WEB), and Cornell Lab of Ornithology (AJM). Thanks to K. Bemis, T. Grande, H. W. Greene, L. Hughes, G. Ortí, L. Page, E. Schuppe, M. Wilson and K. R. Zamudio for discussion and helpful comments on the manuscript. Thanks also to Rick Grosberg for helpful feedback on complexity of aquatic habitats.

## Author Contributions

A.H.B., W.E.B., A.N.R. (listed alphabetically) conceived the study. All authors aggregated data. A.H.B., W.E.B., S.C.F., A.N.R. (listed alphabetically) analyzed the data. S.C.F. and A.N.R. conducted statistical analyses. A.N.R. wrote initial draft; A.H.B., W.E.B., S.C.F., A.N.R. (listed alphabetically) revised, and all authors approved, final version of the manuscript.

## Materials and Correspondence

Requests should be sent to A.N.R. or AHB

## Supplementary Information

**Fig. S1.**
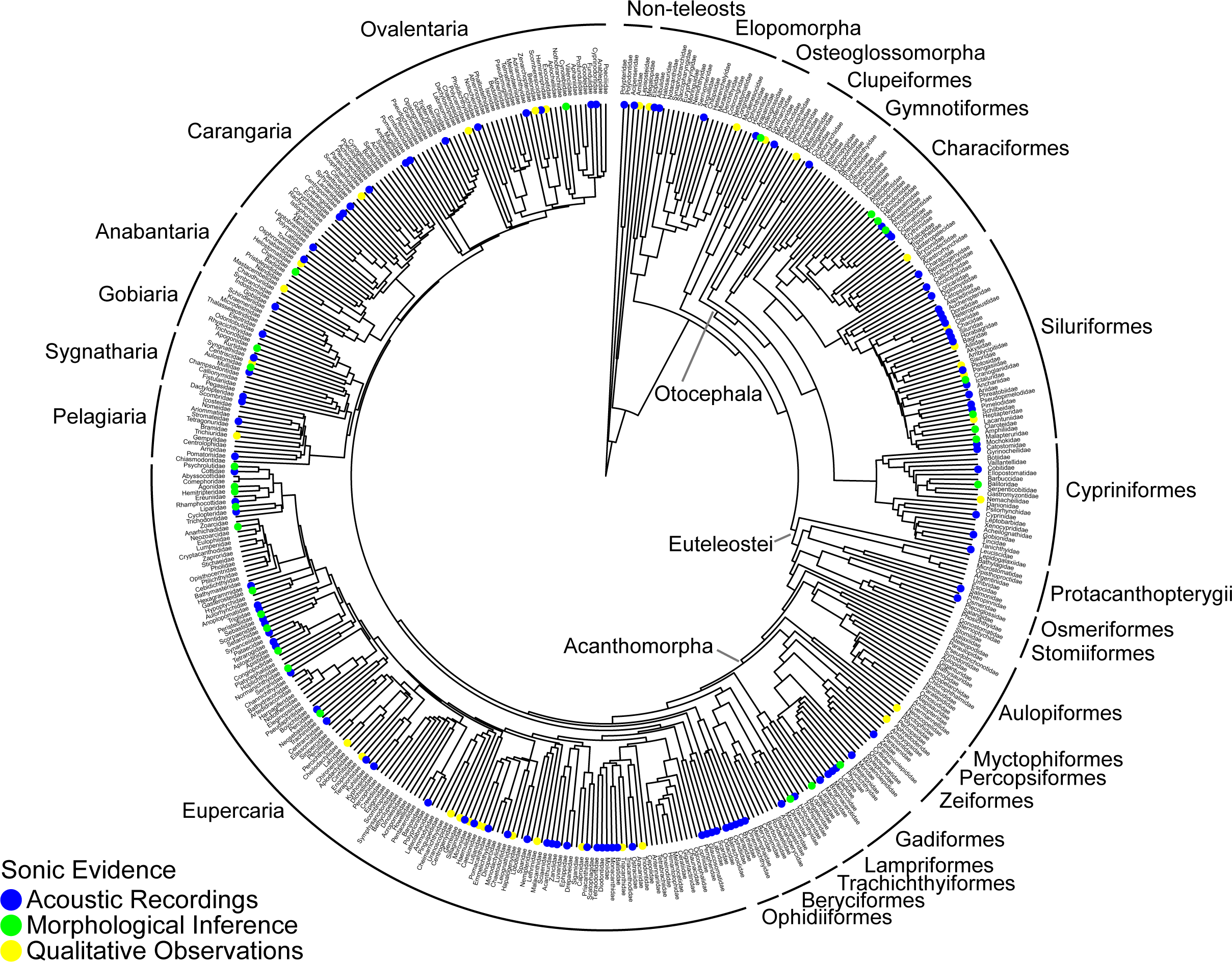
Soniferous behavior mapped onto phylogenetic tree of actinopterygian families. Tree shows three different lines of evidence for soniferous behavior used here and its phylogenetic distribution. Tree is pruned from species-level phylogeny of Rabosky et al. (2018) to family-level here. Some clades recovered using genomic (Betancur-R et al. 2017; Near et al. 2012; Rabosky et al. 2018) and transcriptomic data (Hughes et al. 2018) are supported by well-accepted, anatomical synapomorphies, but others such as Ovalentaria (Hughes et al. 2018) are not.

**Fig. S2.**
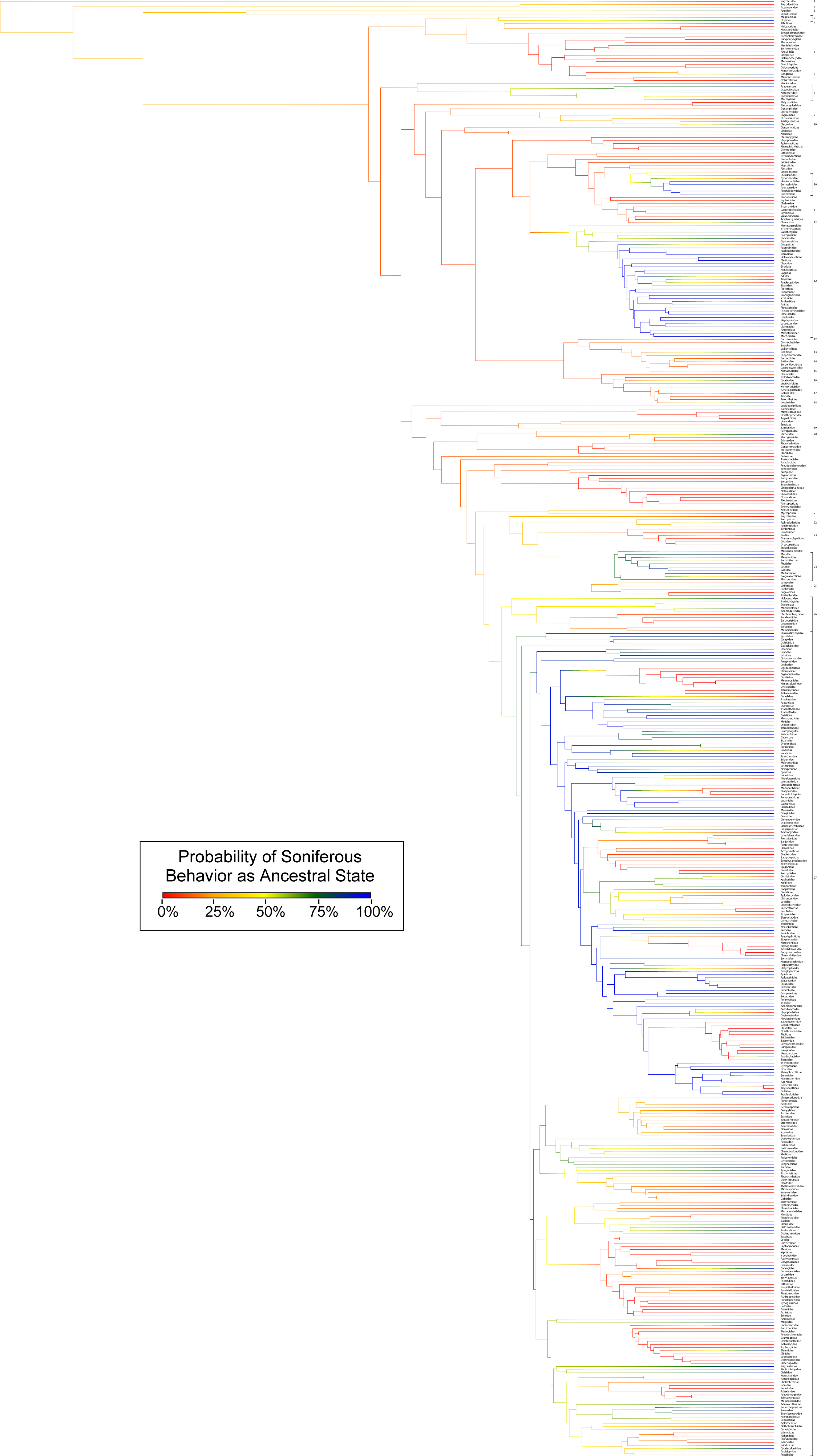
Count of occurrences of the evolution of sonifery. Independent origins of soniferous behavior in actinopterygian fishes, inferred from node values calculated in Fig. 1.

**Fig. S3.**
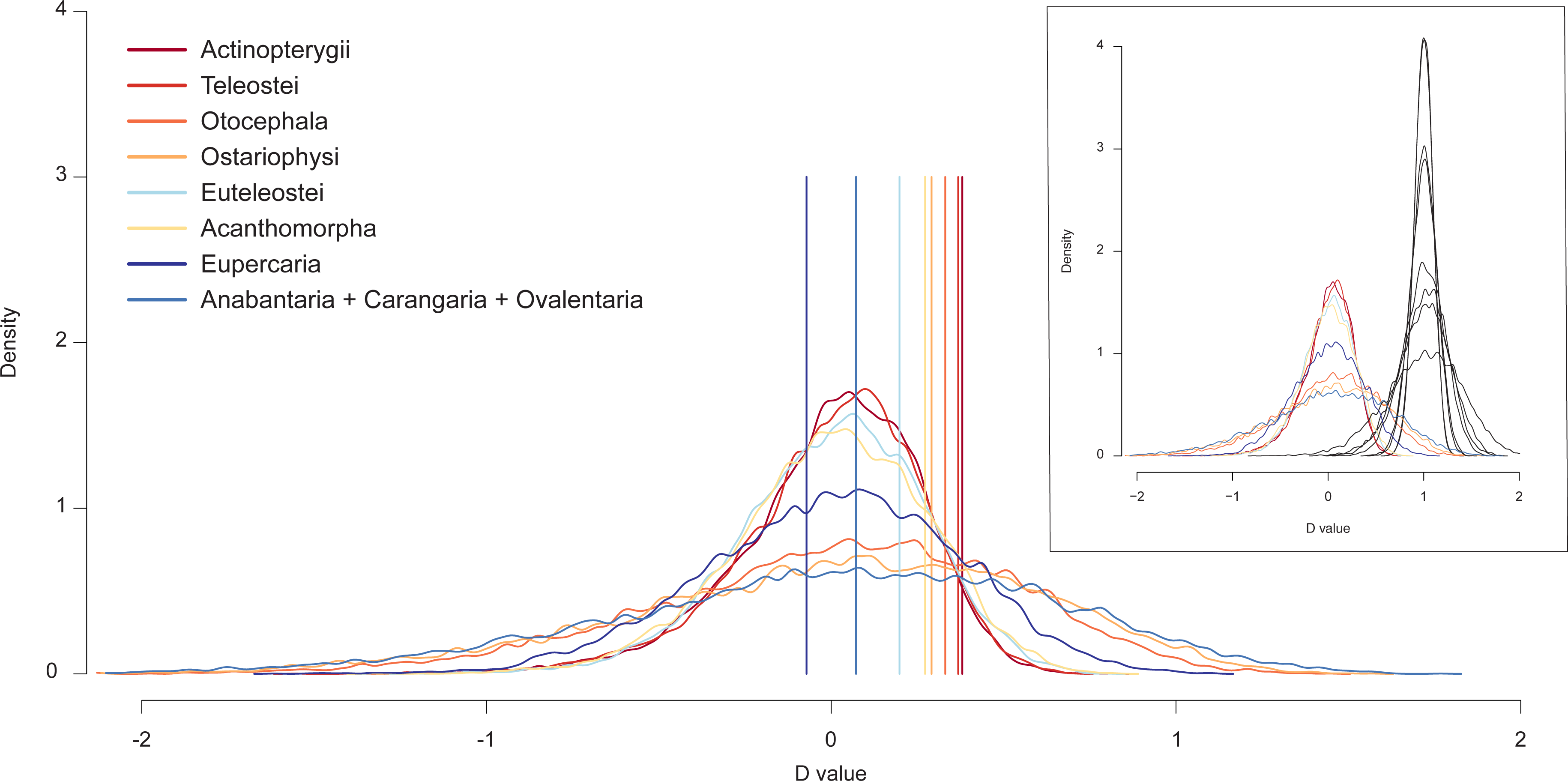
Observed *D*-values for each clade. The observed *D*-values (vertical lines) indicate the strength of phylogenetic signal, based on their value relative to the distribution of simulated *D*-values assuming Brownian evolutionary processes (histograms) for each clade. Values that fall closer to the center of the distribution indicate higher phylogenetic signal within a clade. Some observed *D*-values were closer to (although not near the center of) the simulated distribution based on models of random character evolution with respect to phylogeny (red histograms in upper right plot).

**Table S1.**
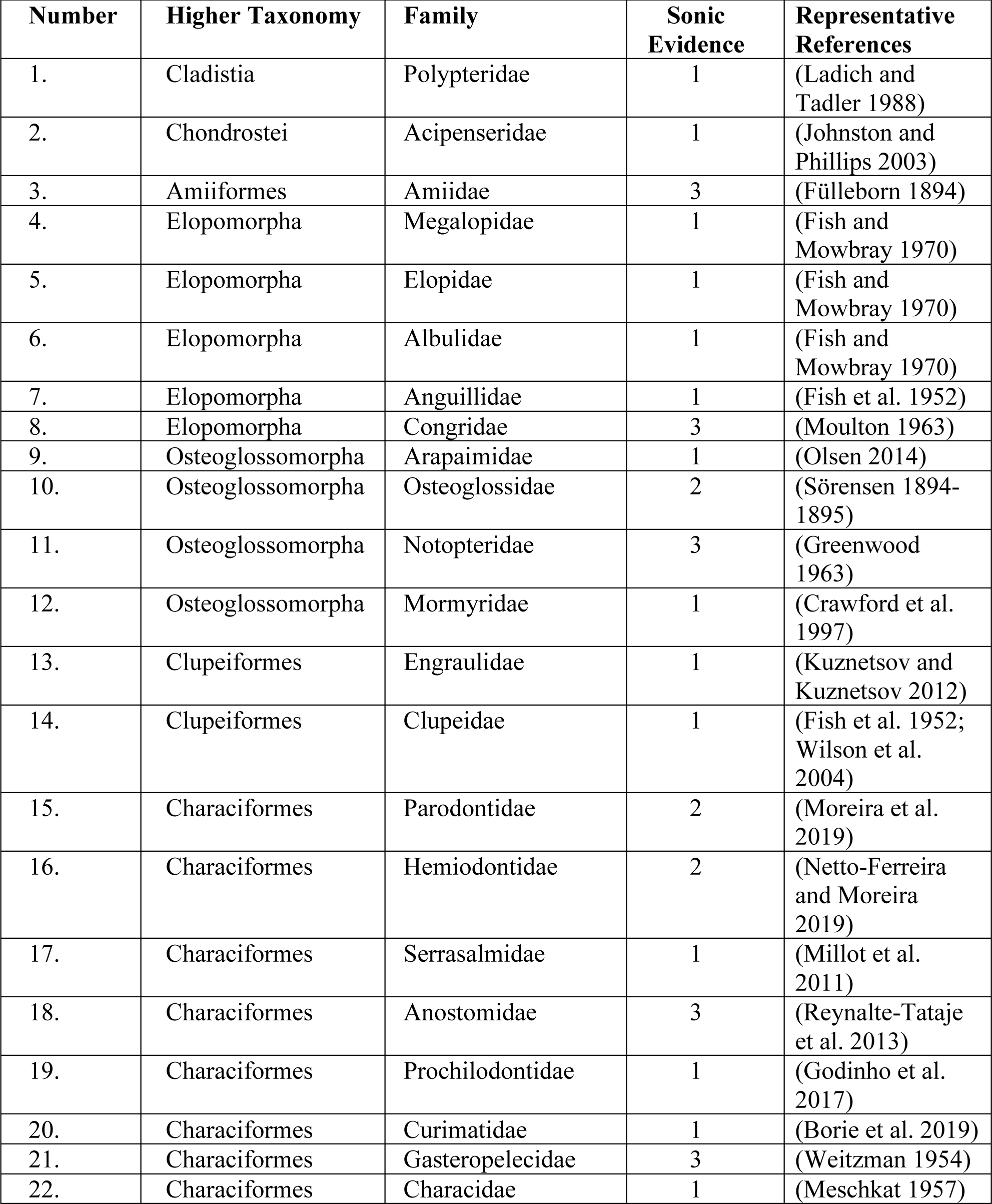
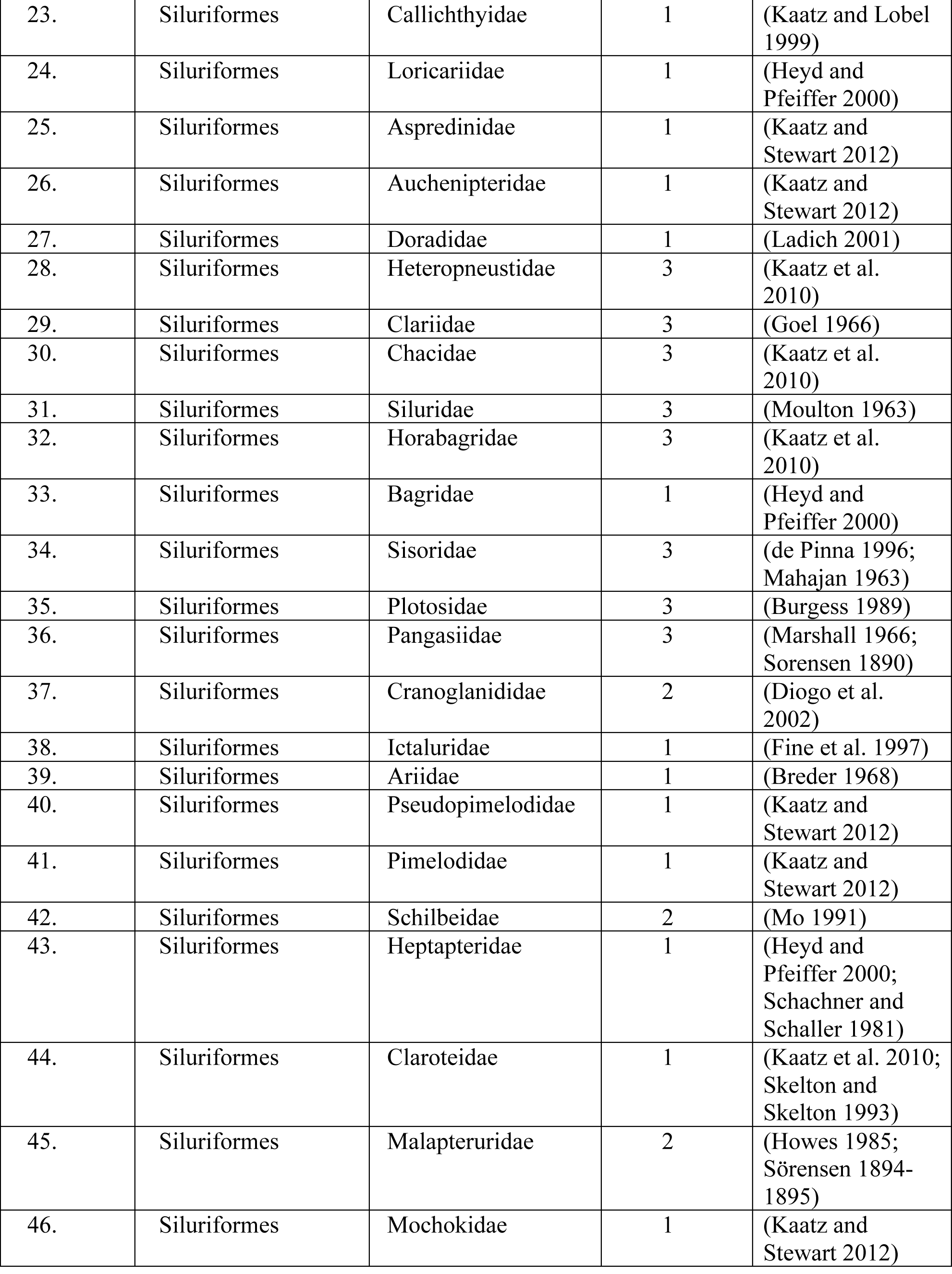
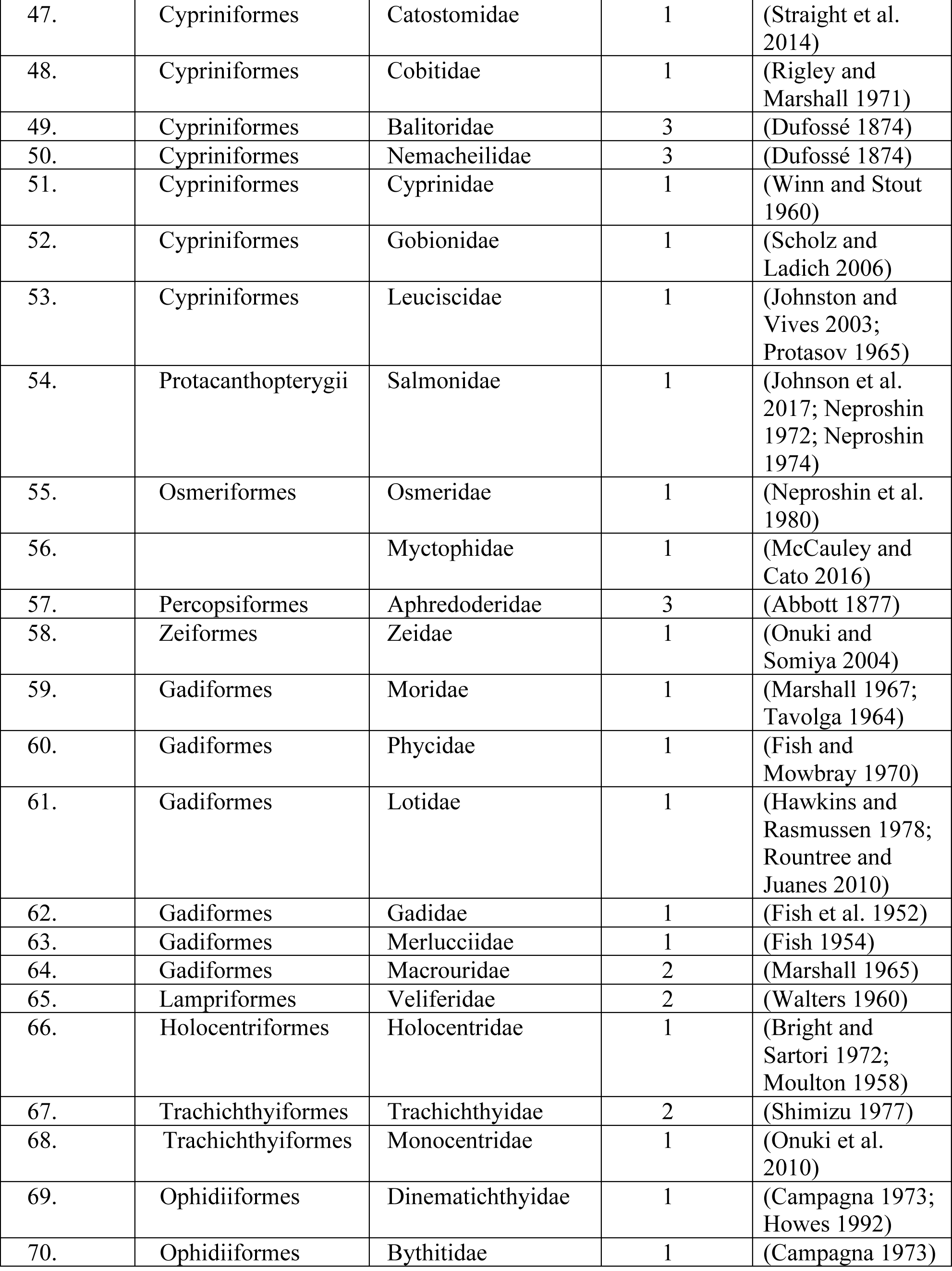
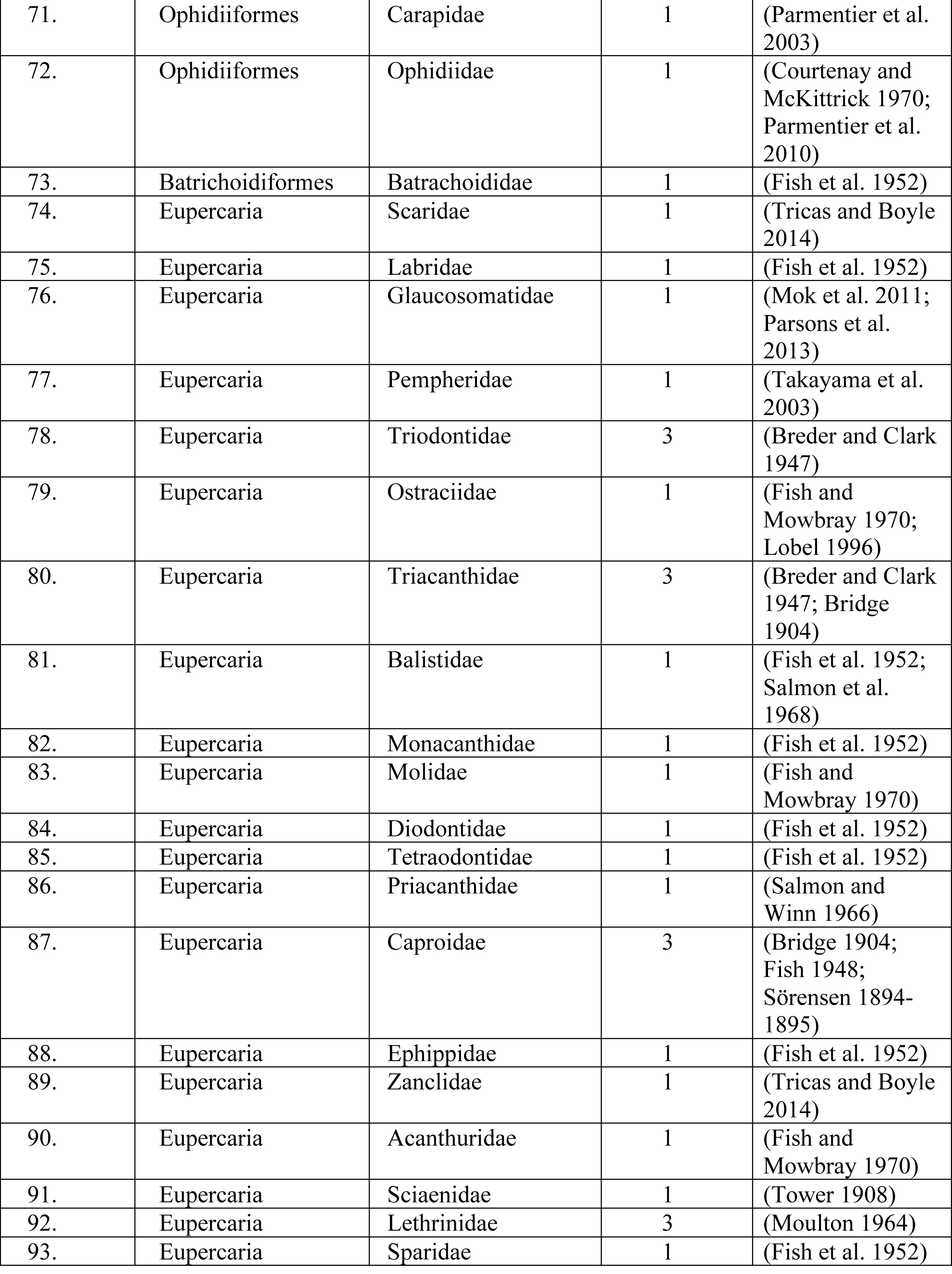
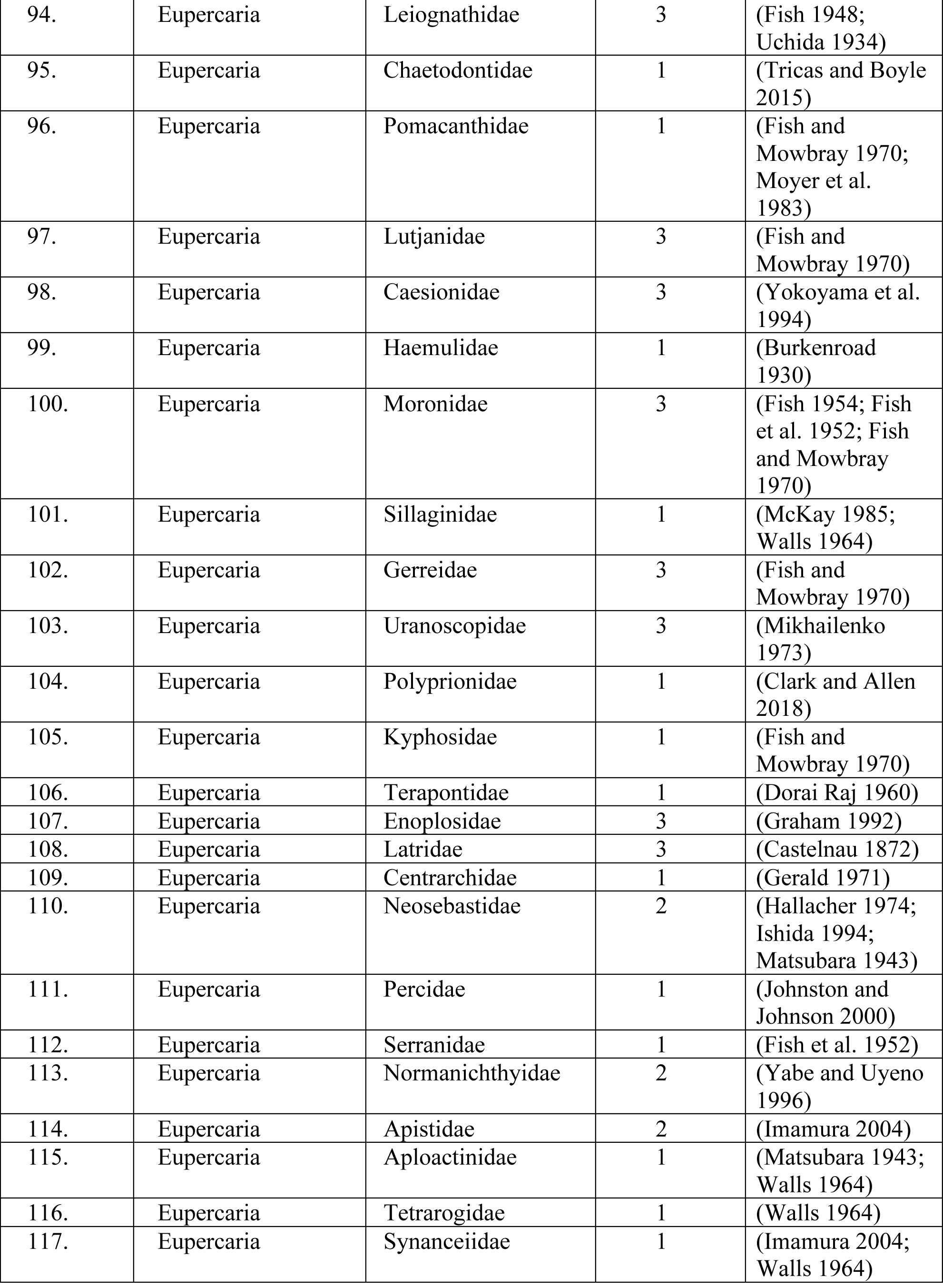
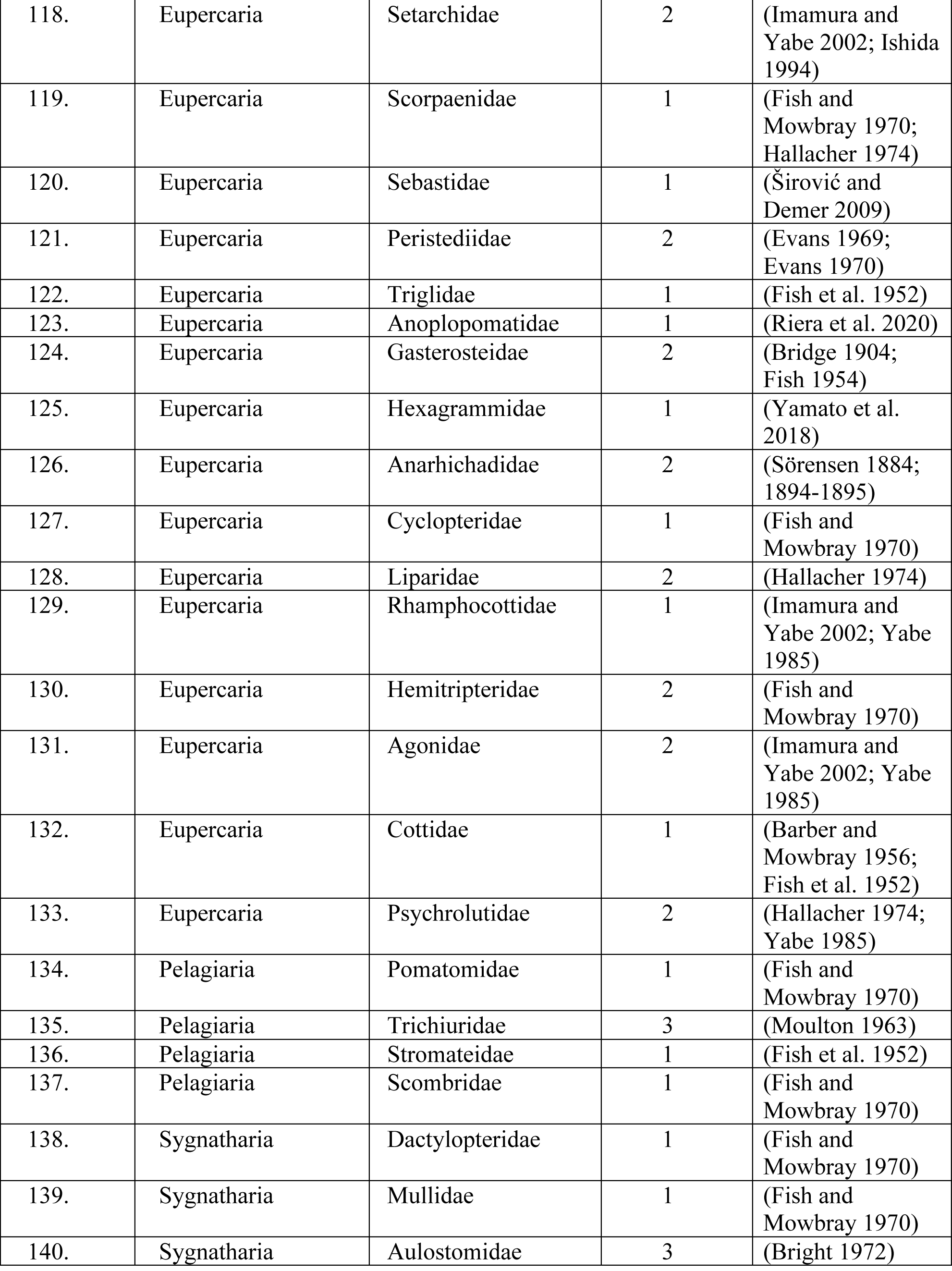
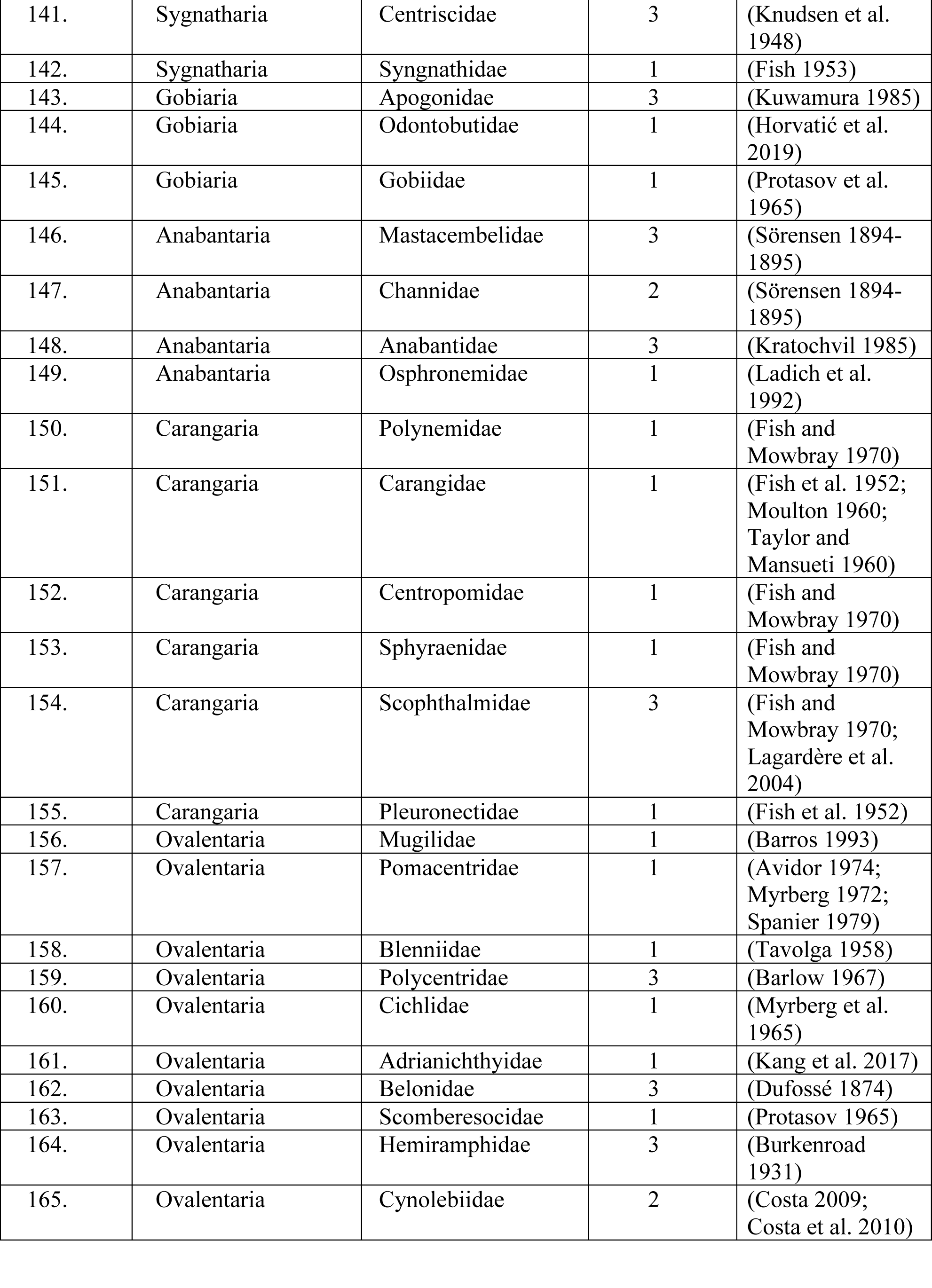
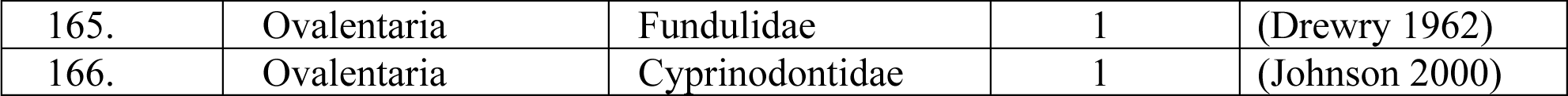
Evidence for sound production in actinopterygian families. Levels of evidence are coded as audio recordings (1), morphological inference (2), or qualitative observations (3). Representative references are included to support evidence of sound production. Families are arranged in sequence following their phylogenetic placement in Figure 1, arranged clockwise.

**Table S2. (separate excel file) Aggregated data for 461 families considered in this analysis.** Showing soniferous behaviors, valid extant species, male alternative reproductive tactics (Mank and Avise 2006), ecological data aggregated from FishBase (Boettiger et al. 2012; Froese and Pauly 2019), including occurrence of families as a function of latitude, salinity, bottom type, habitat, trophic ecology, and reproductive mode.

Nocturnality is strongly correlated with the evolution of acoustic communication within tetrapods (Chen and Wiens 2020). We considered including nocturnality as part of this analysis. Although there are many examples of robust nocturnal chorusing by actinopterygians (Feng and Bass 2016; Mann and Jarvis 2004; McCauley and Cato 2016; Rice et al. 2017), there are no comprehensive analyses of photoperiod-related activity patterns for the soniferous families that are the basis of our study. Assessment of nocturnality in fishes is complicated by potential sampling biases (Dornburg et al. 2017), including seasonal and diel shifts in soniferous and other behaviors coupled to peak times of spawning and reproduction (Feng and Bass 2016; Mann and Jarvis 2004; McCauley and Cato 2016; Rice et al. 2017).

**Table S3.** Clades with independent evolution of sonifery in actinopterygian fishes and associated habitats. Nodes on the phylogenetic tree are labelled in Supplementary Figure 2. Habitat data are from FishBase (Froese and Pauly 2019)

| Clade | Marine/Freshwater | Habitat |
| --- | --- | --- |
| Polypteridae | Freshwater | Demersal |
| Acipenseridae | Anadromous | Demersal spawning and feeding |
| Amiidae | Freshwater | Demersal |
| Megalopidae+Elopidae | Marine | Demersal |
| Albulidae | Marine | Demersal |
| Anguillidae | Anadromous | Demersal in Freshwater |
| Congridae | Marine | Demersal |
| Arapaimidae+Mormyridae+<br>Gymnarchidae+Notopteridae<br>+Osteoglossidae | Freshwater | Rivers |
| Engraulidae | Marine | Some demersal, some pelagic |
| Clupeidae | Marine | Some demersal, some pelagic |
| Curimatoidea | Freshwater | Rivers |
| Gasteropelecidae | Freshwater | Rivers |
| Characidae | Freshwater | Demersal |
| Siluroidei | Mostly Freshwater,<br>some marine | Mostly benthic or demersal |
| Catostomidae | Freshwater | Demersal |
| Cobitidae | Freshwater | Demersal |
| Balitoridae | Freshwater | Demersal |
| Nemacheilidae | Freshwater | Demersal |
| Cyprinidae | Freshwater | Many with demersal spawning and<br>feeding |
| Gobionidae | Freshwater | Rivers, some demersal |
| Leuciscidae | Freshwater | Rivers |
| Salmonidae | Anadromous | Demersal spawning |
| Osmeridae | Anadromous | Demersal and pelagic |
| Myctophidae | Marine | Bathypelagic |
| Aphredoderidae | Freshwater | Demersal spawning |
| Zeidae | Marine | Demersal |
| Gadiformes | Mostly marine | Demersal |

## References

Alfaro, M. E., F. Santini, C. Brock, H. Alamillo, A. Dornburg, D. L. Rabosky, G. Carnevale, L. J. Harmon. 2009. Nine exceptional radiations plus high turnover explain species diversity in jawed vertebrates. Proceedings of the National Academy of Sciences of the United States of America 106:13410–13414.

Amorim, M. C. P., R. O. Vasconcelos, M. Bolgan, S. S. Pedroso, P. J. Fonseca. 2018. Acoustic communication in marine shallow waters: Testing the acoustic adaptive hypothesis in sand gobies. Journal of Experimental Biology 221:jeb183681.

Arcila, D., G. Ortí, R. Vari, J. W. Armbruster, M. L. J. Stiassny, K. D. Ko, M. H. Sabaj, J. Lundberg, L. J. Revell, R. Betancur-R. 2017. Genome-wide interrogation advances resolution of recalcitrant groups in the tree of life. Nature Ecology & Evolution 1:0020.

Bass, A. H., C. W. Clark. 2003. The physical acoustics of underwater sound communication. Pages 15-64 in Acoustic Communication (A. M. Simmons, R. R. Fay, and A. N. Popper, eds.). Springer, New York.

Bass, A. H., G. J. Rose, M. B. Pritz. 2005. Auditory midbrain of fish, amphibians, and reptiles: model systems for understanding auditory function. Pages 459-492 in The Inferior Colliculus (J. A. Winer, and C. E. Schreiner, eds.). Springer, New York.

Bass, A. H. 2014. Central pattern generator for vocalization: is there a vertebrate morphotype? Current Opinion in Neurobiology 28:94–100.

Bass, A. H., B. P. Chagnaud, N. Y. Feng. 2015. Comparative neurobiology of sound production in fishes. Pages 35–75 in Sound Communication in Fishes (F. Ladich, ed.). Springer Vienna, Vienna.

Betancur-R., R., D. Arcila, R. P. Vari, L. C. Hughes, C. Oliveira, M. H. Sabaj, G. Ortí. 2019. Phylogenomic incongruence, hypothesis testing, and taxonomic sampling: The monophyly of characiform fishes. Evolution 73:329–345.

Blomberg, S. P., T. Garland, A. R. Ives. 2003. Testing for phylogenetic signal in comparative data: Behavioral traits are more labile. Evolution 57:717–745.

Blount, Z. D., R. E. Lenski, J. B. Losos. 2018. Contingency and determinism in evolution: Replaying life’s tape. Science 362:eaam5979.

Boettiger, C., D. T. Lang, P. C. Wainwright. 2012. rfishbase: exploring, manipulating and visualizing FishBase data from R. Journal of Fish Biology 81:2030–2039.

Bollback, J. P. 2006. SIMMAP: Stochastic character mapping of discrete traits on phylogenies. BMC Bioinformatics 7:88.

Bose, A. P. H., K. M. Cogliati, N. Luymes, A. H. Bass, M. A. Marchaterre, J. A. Sisneros, B. M. Bolker, S. Balshine. 2018. Phenotypic traits and resource quality as factors affecting male reproductive success in a toadfish. Behavioral Ecology 29:496–507.

Boyle, K. S., O. Colleye, E. Parmentier. 2014. Sound production to electric discharge: Sonic muscle evolution in progress in *Synodontis* spp. catfishes (Mochokidae). Proceedings of the Royal Society B: Biological Sciences 281:20141197.

Bradbury, J. W., S. L. Vehrencamp. 2011. Principles of Animal Communication, 2nd edition. Sinauer Associates, Sunderland, MA.

Braun, C. B., T. Grande. 2008. Evolution of peripheral mechanisms for the enhancement of sound reception. Pages 99-144 in Fish Bioacoustics (J. F. Webb, R. R. Fay, and A. N. Popper, eds.). Springer, New York.

Burns, M. D., D. D. Bloom. 2020. Migratory lineages rapidly evolve larger body sizes than non-migratory relatives in ray-finned fishes. Proceedings of the Royal Society B-Biological Sciences 287:20192615.

Chagnaud, B. P., A. H. Bass. 2013. Vocal corollary discharge communicates call duration to vertebrate auditory system. Journal of Neuroscience 33:18775–18780.

Chen, Z., J. J. Wiens. 2020. The origins of acoustic communication in vertebrates. Nature Communications 11:369.

Colleye, O., B. J. Vetter, R. A. Mohr, L. H. Seeley, J. A. Sisneros. 2019. Sexually dimorphic swim bladder extensions enhance the auditory sensitivity of female plainfin midshipman fish, *Porichthys notatus*. Journal of Experimental Biology 222:jeb204552.

Davis, M. P., N. I. Holcroft, E. O. Wiley, J. S. Sparks, W. L. Smith. 2014. Species-specific bioluminescence facilitates speciation in the deep sea. Marine Biology 161:1139–1148.

Dufossé, M. 1874. Recherches sur les bruits et les sons expressifs que font entendre les poissons d’Europe et sur les organes producteurs de ces phenomenes acoustiques ainsi que sur les appareils de l’audtion de plusieurs de ces animaux. Annales des Sciences Naturelles Cinquième Série: Zoologie et Paléontologie 20:1–134.

Elliott, K. H., C. L. Struik, J. E. Elliott. 2003. Bald eagles, *Haliaeetus leucocephalus*, feeding on spawning plainfin midshipman, *Porichthys notatus*, at Crescent Beach, British Columbia. Canadian Field-Naturalist 117:601–604.

Emlen, S. T., L. W. Oring. 1977. Ecology, sexual selection, and the evolution of mating systems. Science 197:215–223.

Farina, S. C., T. J. Near, W. E. Bemis. 2015. Evolution of the branchiostegal membrane and restricted gill openings in Actinopterygian fishes. Journal of Morphology 276:681–694.

Fine, M. L., E. Parmentier. 2015. Mechanisms of fish sound production. Pages 77–126 in Sound Communication in Fishes (F. Ladich, ed.). Springer Vienna, Vienna.

Fish, M. P., W. H. Mowbray. 1970. Sounds of the Western North Atlantic Fishes. The Johns Hopkins Press, Baltimore.

Forrest, T. G., G. L. Miller, J. R. Zagar. 1993. Sound propagation in shallow water: Implications for acoustic communication by aquatic animals. Bioacoustics 4:259–270.

Fricke, R., W. N. Eschmeyer, J. D. Fong. 2018. Eschmeyer’s Catalog of Fishes: Species by family/Subfamily. Electronic version accessed July 2, 2018. Available at: http://researcharchive.calacademy.org/research/ichthyology/catalog/SpeciesByFamily.asp.

Fricke, R., W. N. Eschmeyer, J. D. Fong. 2020. Eschmeyer’s Catalog of Fishes: Species by family/Subfamily. Electronic version accessed April 4, 2020. Available at: http://researcharchive.calacademy.org/research/ichthyology/catalog/SpeciesByFamily.asp.

Fritz, S. A., A. Purvis. 2010. Selectivity in Mammalian Extinction Risk and Threat Types: a New Measure of Phylogenetic Signal Strength in Binary Traits. Conservation Biology 24:1042–1051.

Froese, R., D. Pauly. 2019. FishBase. Available at: http://www.fishbase.org, version 04/2019.

Giles, S., G.-H. Xu, T. J. Near, M. Friedman. 2017. Early members of ‘living fossil’ lineage imply later origin of modern ray-finned fishes. Nature 549:265–268.

Grosberg, R. K., G. J. Vermeij, P. C. Wainwrigh. 2012. Biodiversity in water and on land. Current Biology 22:R900–R903.

Ho, L. S. T., C. Ané. 2014. A linear-time algorithm for Gaussian and non-Gaussian trait evolution models. Systematic Biology 63:397–408.

Hubbs, C. L. 1920. The bionomics of *Porichthys notatus* Girard. American Naturalist 54:380–384.

Ives, A. R., T. Garland, Jr.,. 2010. Phylogenetic logistic regression for binary dependent variables. Systematic Biology 59:9–26.

Kéver, L., A. H. Bass, E. Parmentier, B. P. Chagnaud. 2020. Neuroanatomical and neurophysiological mechanisms of acoustic and weakly electric signaling in synodontid catfish Journal of Comparative Neurology DOI:10.1002/cne.24920.

Ladich, F., A. Tadler. 1988. Sound production in *Polypterus* (Osteichthyes: Polypteridae). Copeia 1988:1076–1077.

Ladich, F., S. P. Collin, P. Moller, B. G. Kapoor (eds) 2006. Communication in Fishes. Science Publishers, Enfield, N.H.

Ladich, F. 2015. Sound Communication in Fishes. Springer, Vienna.

Ladich, F., H. Winkler. 2017. Acoustic communication in terrestrial and aquatic vertebrates. Journal of Experimental Biology 220:2306–2317.

Lance, M. M., S. J. Jeffries. 2009. Harbor seal diet in Hood Canal, South Puget Sound and the San Juan Island Archipelago. Contract Report to Pacific States Marine Fisheries Commission for Job Code 497; NOAA Award No. NA05NMF4391151. Washington Department of Fish and Wildlife, Olympia, WA.

Lee, J. S. F., A. H. Bass. 2006. Dimorphic male midshipman fish: reduced sexual selection or sexual selection for reduced characters? Behavioral Ecology 17:670–675.

Lugli, M. 2015. Habitat acoustics and the low-frequency communication of shallow water fishes. Pages 175–206 in Sound Communication in Fishes (F. Ladich, ed.). Springer, Vienna.

Mank, J. E., J. C. Avise. 2006. Comparative phylogenetic analysis of male alternative reproductive tactics in ray-finned fishes. Evolution 60:1311–1316.

Mann, D., J. Locascio, C. Wall. 2016. Listening in the ocean: New discoveries and insights on marine life from autonomous passive acoustic recorders. Pages 309–324 in Listening in the Ocean (W. W. L. Au, and M. O. Lammers, eds.). Springer New York, New York, NY.

McCabe, E. J. B., D. P. Gannon, N. B. Barros, R. S. Wells. 2010. Prey selection by resident common bottlenose dolphins (*Tursiops truncatus*) in Sarasota Bay, Florida. Marine Biology 157:931–942.

Miles, M. C., F. Goller, M. J. Fuxjager. 2018. Physiological constraint on acrobatic courtship behavior underlies rapid sympatric speciation in bearded manakins. Elife 7:e40630.

Miles, M. C., M. J. Fuxjager. 2019. Phenotypic diversity arises from secondary signal loss in the elaborate visual displays of toucans and barbets. American Naturalist 194:152–167.

Myrberg, A. A., R. J. Riggio. 1985. Acoustically mediated individual recognition by a coral reef fish (*Pomacentrus partitus*). Animal Behaviour 33:411–416.

Navia, A. F., P. A. Majia-Fola, A. Giraldo. 2007. Feeding ecology of elasmobranchs in coastal waters of the Colombian Eastern Tropical Pacific. BMC Ecology 7:8.

Nelson, J. S., T. C. Grande, M. V. H. Wilson. 2016. Fishes of the World, 5th Edition. John Wiley & Sons, Inc., Hoboken, NJ.

Orme, D., R. Freckleton, G. Thomas, T. Petzoldt, S. Fritz, N. Isaac, W. Pearse. 2013. *caper: Comparative analyses of phylogenetics and evolution in R*. R package version 0.5.2. https://CRAN.R-project.org/package=caper.

Popper, A. N., B. M. Casper. 2011. Fish bioacoustics: an Introduction. Pages 236-243 in Encyclopedia of Fish Physiology (A. P. Farrell, ed.). Academic Press, San Diego.

Rabosky, D. L., J. Chang, P. O. Title, P. F. Cowman, L. Sallan, M. Friedman, K. Kaschner, C. Garilao, T. J. Near, M. Coll, M. E. Alfaro. 2018. An inverse latitudinal gradient in speciation rate for marine fishes. Nature 559:392–395.

Radford, C. A., J. C. Montgomery, P. Caiger, P. Johnston, J. Lu, D. M. Higgs. 2013. A novel hearing specialization in the New Zealand bigeye, *Pempheris adspersa*. Biology Letters 9:4.

Revell, L. J. 2012. phytools: an R package for phylogenetic comparative biology (and other things). Methods in Ecology and Evolution 3:217–223.

Rice, W. R. 1989. Analyzing tables of statistical tests. Evolution 43:223–225.

Rohmann, K. N., D. Fergus, A. H. Bass. 2013. Plasticity in ion channel expression underlies variation in hearing during reproductive cycles. Current Biology 23:678–683.

Smith, W. L., W. C. Wheeler. 2006. Venom evolution widespread in fishes: a phylogenetic road map for the bioprospecting of piscine venoms. Journal of Heredity 97:206–217.

Stewart, T. A., W. L. Smith, M. I. Coates. 2014. The origins of adipose fins: an analysis of homoplasy and the serial homology of vertebrate appendages. Proceedings of the Royal Society B: Biological Sciences 281:20133120.

Tower, R. W. 1908. The production of sound in the drumfishes, the sea-robin and the toadfish. Annals of the New York Academy of Sciences 18:149–180.

Urick, R. J. 1975. Principles of Underwater Sound, 2nd edition. McGraw-Hill, New York.

von Frisch, K. 1938. The sense of hearing in fish. Nature 141:8–11.

Wainwright, P. C., S. J. Longo. 2017. Functional innovations and the conquest of the oceans by acanthomorph fishes. Current Biology 27:R550–R557.

Ward, A. B., E. B. Brainerd. 2007. Evolution of axial patterning in elongate fishes. Biological Journal of the Linnean Society 90:97–116.

Wilkins, M. R., N. Seddon, R. J. Safran. 2013. Evolutionary divergence in acoustic signals: causes and consequences. Trends in Ecology & Evolution 28:156–166.

Zhang, Y. S., A. A. Ghazanfar. 2020. A hierarchy of autonomous systems for vocal production. Trends in Neurosciences 43:115–126.

## Supplementary References

Abbott CC. 1877. Traces of a voice in Fishes. American Naturalist 11:147–156.

Avidor A. 1974. The signal jump and its associated sound in fish of the genus Dascyllus from the Gulf of Eilat. Master’s Thesis. Tel Aviv University.

Barber SB, Mowbray WH. 1956. Mechanism of sound production in the sculpin. Science 124:219–220.

Barlow GW. 1967. Social behavior of a South American leaf fish *Polycentrus schomburgkii* with an account of recurring pseudofemale behavior. American Midland Naturalist 78:215–234.

Barros NB. 1993. Feeding ecology and foraging strategies of bottlenose dolphins on the central east coast of Florida. Coral Gables, FL: University of Miami.

Betancur-R R, Wiley EO, Arratia G, Acero A, Bailly N, Miya M, Lecointre G, Ortí G. 2017. Phylogenetic classification of bony fishes. BMC Evolutionary Biology 17:162.

Boettiger C, Lang DT, Wainwright PC. 2012. rfishbase: Exploring, manipulating and visualizing FishBase data from R. Journal of Fish Biology 81:2030–2039.

Borie A, Hungria DB, Ali H, Doria CR, Fine ML, Travassos PE. 2019. Disturbance calls of five migratory Characiformes species and advertisement choruses in Amazon spawning sites. Journal of Fish Biology 95:820–832.

Breder CM, Clark E. 1947. A contribution to the visceral anatomy, development and relationships of the Plectognathi. Bulletin of the American Museum of Natural History 88:287–319.

Breder CM, Jr. 1968. Seasonal and diurnal occurrences of fish sounds in a small Florida bay. Bulletin of the American Museum of Natural History 138:327–378.

Bridge TW. 1904. Fishes. Cambridge Natural History vii:pp. 139–537.

Bright CM. 1972. Bio-acoustic studies on reef organisms. In: Collette BB, Earle SA, editors. Results of the Tektite Program: Ecology of Coral Reef Fishes. Los Angeles: Natural History Museum, Los Angeles County, Science Bulletin 14. p. 45–69.

Bright TJ, Sartori JD. 1972. Sound production by the reef fishes *Holocentrus coruscus*, *Holocentrus rufus* and *Myripristis jacobus*, Family Holocentridae. Hydro-Lab Journal 1:11–20.

Burgess WE. 1989. An atlas of freshwater and marine catfishes: a preliminary survey of the Siluriformes. Neptune CIty: T. F. H. Publications, Inc.

Burkenroad MD. 1930. Sound production in the Haemulidae. Copeia 1930:17–18.

Burkenroad MD. 1931. Notes on the sound-producing marine fishes of Louisiana. Copeia 1931:20–28.

Campagna A. 1973. Comparative morphology and phylogenetic significance of sound-producing mechanisms in some Brotulidae (Osteichthyes, Ophidioidea). Doctoral Thesis. Boston University.

Castelnau FL. 1872. Contribution to the ichthyology of Australia. Proceedings of the Zoological and Acclimitisation Society of Victoria 1:29–247.

Chen Z, Wiens JJ. 2020. The origins of acoustic communication in vertebrates. Nature Communications 11:369.

Clark BLF, Allen LG. 2018. Field observations on courtship and spawning behavior of the giant sea bass, *Stereolepis gigas*. Copeia 106:171–179.

Costa WJEM. 2009. Morphology of the teleost pharyngeal jaw apparatus in the Neotropical annual killifish genus Cynolebias (Cyprinodontiformes: Aplocheiloidei: Rivulidae). Cybium 33:145–150.

Costa WJEM, Ramos TPA, Alexandre LC, Ramos RTC. 2010. *Cynolebias parnaibensis*, a new seasonal killifish from the Caatinga, Parnaíba River basin, northeastern Brazil, with notes on sound producing courtship behavior (Cyprinodontiformes: Rivulidae). Neotropical Ichthyology 8:283–288.

Courtenay WR, Jr., McKittrick FA. 1970. Sound-producing mechanisms in carapid fishes, with notes on phylogenetic implications. Marine Biology 7:131–137.

Crawford JD, Cook AP, Herberlein AS. 1997. Bioacoustic behavior of African fishes (Mormyridae):potential cues for species and individual recognition in *Pollimyrus*. Journal of the Acoustical Society of America 102:1200–1212.

de Pinna MCC. 1996. A phylogenetic analysis of the Asian catfish families Sisoridae, Akysidae, and Amblycipitidae, with a hypothesis on the relationships of the neotropical Aspredinidae (Teleostei, Ostariophysi). Fieldiana Zoology 84:1–83.

Diogo R, Chardon M, Vandewalle P. 2002. Osteology and myology of the cephalic region and pectoral girdle of the Chinese catfish *Cranoglanis bouderius*, with a discussion on the autapomorphies and phylogenetic relationships of the Cranoglanididae (Teleostei : Siluriformes). Journal of Morphology 253:229–242.

Dorai Raj BS. 1960. On the production of underwater sound by *Therapon jarbua*. Current Science 29:277–278.

Dornburg A, Forrestel EJ, Moore JA, Iglesias TL, Jones A, Rao L, Warren DL. 2017. An assessment of sampling biases across studies of diel activity patterns in marine ray-finned fishes (Actinopterygii). Bulletin of Marine Science 93:611–639.

Drewry GE. 1962. Some observations of courtship behavior and sound production in five species of Fundulus Masters Thesis. Austin: University of Texas.

Dufossé M. 1874. Recherches sur les bruits et les sons expressifs que font entendre les poissons d’Europe et sur les organes producteurs de ces phenomenes acoustiques ainsi que sur les appareils de l’audtion de plusieurs de ces animaux. Annales des Sciences Naturelles Cinquième Série: Zoologie et Paléontologie 20:1–134.

Evans RR. 1969. Phylogenetic significance of sound producing mechanisms of Western Atlantic fishes of the family Triglidae and Peristediidae Doctoral Thesis. Ann Arbor, Michigan: Boston University Graduate School.

Evans RR. 1970. Phylogenetic significance of teleost sound producing mechanisms. Journal of the Colorado-Wyoming Academy of Science 7:9–10.

Feng NY, Bass AH. 2016. “Singing” fish rely on circadian rhythm and melatonin for the timing of nocturnal courtship vocalization. Current Biology 26:2681–2689.

Fine ML, Friel JP, McElroy D, King CB, Loesser KE, Newton S. 1997. Pectoral spine locking and sound production in the channel catfish, *Ictalurus punctatus*. Copeia 1997:777–790.

Fish MP. 1948. Sonic fishes of the Pacific. Woods Hole, MA: Woods Hole Oceanographic Institution.

Fish MP. 1953. The production of underwater sound by the northern seahorse, *Hippocampus hudsonius*. Copeia 1953:98–99.

Fish MP. 1954. The character and significance of sound production among fishes of the Western North Atlantic. Bulletin of the Bingham Oceanographic Collection 14:1–109.

Fish MP, Kelsey AS, Mowbray WH. 1952. Studies on the production of underwater sound by North Atlantic coastal fishes. Journal of Marine Research 11:180–193.

Fish MP, Mowbray WH. 1970. Sounds of the Western North Atlantic Fishes. Baltimore: The Johns Hopkins Press.

Froese R, Pauly D. 2019. FishBase. Available at: http://www.fishbase.org, version 04/2019.

Fülleborn F. 1894. Bericht über eine zur Untersuchung der Entwickelung von Amia, Lepidosteus und Necturus unternommene Reise nach Nord-America. Sitzungsberichte Deutche Akademie der Wissenschaften zu Berlin 40:1057–1070.

Gerald JW. 1971. Sound production during courtship in six species of sunfish (Centrarchidae). Evolution 25:75–87.

Godinho AL, Silva CCF, Kynard B. 2017. Spawning calls by zulega, *Prochilodus argenteus*, a Brazilian riverine fish. Environmental Biology of Fishes 100:519–533.

Goel HC. 1966. Sound production in *Clarias batrachus* (Linnaeus). Copeia 1966:622–624.

Graham R. 1992. Sounds fishy. Australia’s Geographic Magazine 14:76–83.

Greenwood PH. 1963. The swimbladder in African Notopteridae (Pisces) and its bearing on the taxonomy of the family. Bulletin of the British Museum of Natural History: Zoology 11:377–412.

Hallacher LE. 1974. The comparative morphology of extrinsic gasbladder musculature in the scorpionfish genus *Sebastes* (Pisces: Scorpaenidae). Proceedings of the California Academy of Sciences 40:59–86.

Hawkins AD, Rasmussen KJ. 1978. The calls of gadoid fish. Journal of the Marine Biological Association of the UK 58:891–911.

Heyd A, Pfeiffer W. 2000. Über die Lauterzeugung der Welse (Siluroidei, Ostariophysi, Teleostei) und ihren Zusammenhang mit der Phylogenie und der schreckreaktion. Revue Suisse de Zoologie 107:165–211.

Horvatić S, Malavasi S, Parmentier E, Marčić Z, Buj I, Mustafić P, Ćaleta M, Smederevac-Lalić M, Skorić S, Zanella D. 2019. Acoustic communication during reproduction in the basal gobioid Amur sleeper and the putative sound production mechanism. Journal of Zoology DOI:10.1111/jzo.12719.

Howes GJ. 1985. The phylogenetic relationships of the electric catfish family Malapteruridae (Teleostei, Silurodei). Journal of Natural History 19:37–67.

Howes GJ. 1992. Notes on the anatomy and classification of ophidiiform fishes with particular reference to the abyssal genus *Acanthonus* Günther, 1878. Bulletin of the British Museum of Natural History: Zoology 58:95–131.

Hughes LC, Ortí G, Huang Y, Sun Y, Baldwin CC, Thompson AW, Arcila D, Betancur-R. R, Li C, Becker L, Bellora N, Zhao X, Li X, Wang M, Fang C, Xie B, Zhou Z, Huang H, Chen S, Venkatesh B, Shi Q. 2018. Comprehensive phylogeny of ray-finned fishes (Actinopterygii) based on transcriptomic and genomic data. Proceedings of the National Academy of Sciences of the United States of America.

Imamura H. 2004. Phylogenetic relationships and new classification of the superfamily Scorpaenoidea (Actinopterygii: Perciformes). Species Diversity 9:1–36.

Imamura H, Yabe M. 2002. Demise of the scorpaeniformes (Actinopterygii: Percomorpha): An alternative phylogenetic hypothesis. Bulletin of Fisheries Sciences Hokkaido University 53:107–128.

Ishida M. 1994. Phylogeny of the suborder Scorpaenoidei (Pisces: Scorpaeniformes). Bulletin of Nansei National Fisheries Research Institute 27:1–112.

Johnson DL. 2000. Sound Production in *Cyprinodon bifasciatus* (Cyprinodontiformes). Environmental Biology of Fishes 59:341–346.

Johnson NS, Higgs D, Binder TR, Marsden JE, Buchinger T, Brege L, Bruning T, Farha S, Krueger CC. 2017. Evidence of sound production by spawning lake trout (*Salvelinus namaycush*) in lakes Huron and Champlain. Canadian Journal of Fisheries and Aquatic Sciences 75:429–438.

Johnston CE, Johnson DL. 2000. Sound production during the spawning season in cavity-nesting darters of the subgenus *Catonotus* (Percidae : Etheostoma). Copeia 2000:475–481.

Johnston CE, Phillips CT. 2003. Sound production in sturgeon *Scaphirhynchus albus* and *S. platorynchus* (Acipenseridae). Environmental Biology of Fishes 69:59–64.

Johnston CE, Vives SP. 2003. Sound production in *Codoma ornata* (Girard) (Cyprinidae). Environmental Biology of Fishes 68:81–85.

Kaatz IM, Lobel PS. 1999. Acoustic behavior and reproduction in five species of *Corydoras* catfishes (Callichthyidae). Biological Bulletin 197:242–242.

Kaatz IM, Stewart DJ. 2012. Bioacoustic variation of swimbladder disturbance sounds in Neotropical doradoid catfishes (Siluriformes: Doradidae, Auchenipteridae): potential morphological correlates. Current Zoology 58:171–188.

Kaatz IM, Stewart DJ, Rice AN, Lobel PS. 2010. Differences in pectoral fin spine morphology between vocal and silent clades of catfishes (Order Siluriformes): ecomorphological implications. Current Zoology 56:73–89.

Kang IJ, Qiu X, Moroishi J, Oshima Y. 2017. Sound production in Japanese medaka (*Oryzias latipes*) and its alteration by exposure to aldicarb and copper sulfate. Chemosphere 181:530–535.

Knudsen VO, Alford RS, Emling JW. 1948. Underwater ambient noise. Journal of Marine Research 7:410–429.

Kratochvil H. 1985. Beitraege zur Lautiologi der Anabantodei: Bau, Funktion und Entwicklung von lauterzeugenden Systemen. Zoologische Jahrbücher Abteilung für Allegmeine Zoologie und Physiologie der Tiere 89:203–255.

Kuwamura T. 1985. Social and reproductive behavior of three mouthbrooding cardinalfishes, *Apogon doederleini*, A. niger and A. notatus. Environmental Biology of Fishes 13:17–24.

Kuznetsov MY, Kuznetsov YA. 2012. Sound production in some physostomous fish species and effects of biological sounds on fish. In: Popper AN, Hawkins A, editors. Effects of Noise on Aquatic Life. New York: Springer. p. 177–180.

Ladich F. 2001. Sound-generating and -detecting motor system in catfish: design of swimbladder muscles in doradids and pimelodids. Anatomical Record 263:297–306.

Ladich F, Bischof C, Schleinzer G, Fuchs A. 1992. Intra- and interspecific differences in agonistic vocalizations in croaking Gouramis (Genus: *Trichopsis*, Anabantoidei, Teleostei). Bioacoustics 4:131–141.

Ladich F, Tadler A. 1988. Sound production in *Polypterus* (Osteichthyes: Polypteridae). Copeia 1988:1076–1077.

Lagardère JP, Mallekh R, Mariani A. 2004. Acoustic characteristics of two feeding modes used by brown trout (*Salmo trutta*), rainbow trout (*Oncorhynchus mykiss*) and turbot (*Scophthalmus maximus*). Aquaculture 240:607–616.

Lobel PS. 1996. Spawning sound of the trunkfish, *Ostracion meleagris* (Ostraciidae). Biological Bulletin 191:308–309.

Mahajan CL. 1963. Sound producing apparatus in an Indian catfish *Sisor rhabdophorus* Hamilton. Journal of Linnean Society 44:721–724.

Mank JE, Avise JC. 2006. Comparative phylogenetic analysis of male alternative reproductive tactics in ray-finned fishes. Evolution 60:1311–1316.

Mann DA, Jarvis SM. 2004. Potential sound production by a deep-sea fish. Journal of the Acoustical Society of America 115:2331–2333.

Marshall NB. 1965. Systematic and biological studies of the Macrourid fishes (Anacanthini-Teleostii). Deep-Sea Research 12:299–322.

Marshall NB. 1966. The Life of Fishes. Cleveland: The World Publishing Company.

Marshall NB. 1967. Sound-producing mechanisms and the biology of deep-sea fishes. In: Tavolga WN, editor. Marine Bioacoustics. Oxford: Pergamon Press. p. 123–133.

Matsubara K. 1943. Studies on the scorpaenoid fishes of Japan: anatomy, phylogeny and taxonomy. Trans Sigenkagaku Kenkyusho 1:1–170.

McCauley RD, Cato DH. 2016. Evening choruses in the Perth Canyon and their potential link with Myctophidae fishes. Journal of the Acoustical Society of America 140:2384–2398.

McKay RJ. 1985. A revision of the fishes of the family Sillaginidae. Memoirs of the Queensland Museum 22:1–74.

Meschkat A. 1957. Von den Stimmen der Fishe im Amazonas. Fischwirt Zeitschrift für Seen-und Flussfischen 3:67–68.

Mikhailenko NA. 1973. Organ of sound formation and electro generation in the Black Sea stargazer *Uranoscopus scaber* (Uranoscopidae). Zoologicheskiĭ Zhurnal 52:1353–1359.

Millot S, Vandewalle P, Parmentier E. 2011. Sound production in red-bellied piranhas (*Pygocentrus nattereri*, Kner): an acoustical, behavioral and morphofunctional study. Journal of Experimental Biology 214:3613–3618.

Mo T. 1991. Anatomy, relationships and systematics of the Bagridae (Teleostei: Siluroidei) with a hypothesis of siluroid phylogeny. Theses Zoologicae 17:1–216.

Mok H-K, Parmentier E, Chiu K-H, Tsai K-E, Chiu P-H, Fine M. 2011. An intermediate in the evolution of superfast sonic muscles. Frontiers in Zoology 8:31.

Moreira CR, Netto-Ferreira AL, Colaco MV, Nogueira LP. 2019. How parodontid sishes (Ostariophysi: Characiformes) sing with their ribs. Journal of Morphology 280:S185.

Moulton JM. 1958. The acoustical behavior of some fishes in the Bimini area. Biological Bulletin 114:357–374.

Moulton JM. 1960. The acoustical anatomy of teleost fishes. Anatomical Record 138:371–372.

Moulton JM. 1963. Acoustic behavior of fishes. In: Busnel RG, editor. Acoustic behavior of animals. Amsterdam: Elsevier. p. 655–685.

Moulton JM. 1964. Acoustic behavior of fishes. In: Busnel RG, editor. Acoustic behavior of animals. New York: Elsevier. p. 655–693.

Moyer JT, Thresher RE, Collin PL. 1983. Courtship, spawning and inferred social organization of American angelfishes (genera *Pomacanthus*, Holacanthus and Centropyge; Pomacanthidae). Environmental Biology of Fishes 9:25–39.

Myrberg AA, Jr. 1972. Ethology of the bicolor damselfish, Eupomacentrus partitus (Pisces: Pomacentridae): a comparison of laboratory and field behavior. Animal Behavior Monographs 5:199–283.

Myrberg AA, Jr., Kramer E, Heinecke P. 1965. Sounds produced by cichlid fishes. Science 149:555–558.

Near TJ, Eytan RI, Dornburg A, Kuhn KL, Moore JA, Davis MP, Wainwright PC, Friedman M, Smith WL. 2012. Resolution of ray-finned fish phylogeny and timing of diversification. Proceedings of the National Academy of Sciences of the United States of America 109:13698–13703.

Neproshin AY. 1972. Some physical characteristics of the sounds in Pacific salmon. Zoologicheskiĭ Zhurnal 51:1025–1030.

Neproshin AY. 1974. The acoustic behavior of some far eastern salmon in the spawning period. Journal of Ichthyology 14:154–157.

Neproshin AY, Kruchinin ON, Fedoseenkov AS. 1980. The acoustical behavior of the pond smelt, Hypomesus olidus. Journal of Ichthyology 19:167–169.

Netto-Ferreira AL, Moreira CR. 2019. A review of the vocal species of characiformes (Teleostei: Ostariophysi). Journal of Morphology 280:S191.

Olsen JEB. 2014. Predation Avoidance Mechanisms of Juvenile Arapaima spp.: Significance of Synchronized Breathing and Sound Production. State University of New York College of Environmental Science and Forestry.

Onuki A, Somiya H. 2004. Two types of sounds and additional spinal nerve innervation to the sonic muscle in John Dory, *Zeus faber* (Zeiformes: Teleostei). Journal of the Marine Biological Association of the UK 84:843–850.

Onuki A, Takizawa T, Yamamoto N, Somiya H. 2010. Sound characteristics and sonic motor system in the pineconefish, *Monocentris japonica* (Beryciformes: Monocentridae). Copeia 2010:531–539.

Parmentier E, Bouillac G, Dragicevic B, Dulcic J, Fine M. 2010. Call properties and morphology of the sound-producing organ in *Ophidion rochei* (Ophidiidae). Journal of Experimental Biology 213:3230–3236.

Parmentier E, Vandewalle P, Lagardère JP. 2003. Sound-producing mechanisms and recordings in Carapini species (Teleostei, Pisces). Journal of Comparative Physiology A 189:283–292.

Parsons M, McCauley R, Thomas F. 2013. The sounds of fish off Cape Naturaliste, Western Australia. Acoustics Australia 41:58–64.

Protasov VR. 1965. Bioacoustics of Fishes. Moscow: Nauka.

Protasov VR, Tzvetkov VI, Rascheperin VK. 1965. Acoustic signals emitted by *Neogobius melanostomus* Pallas. Zhurnal Obshchei Biologii 26:151–160.

Rabosky DL, Chang J, Title PO, Cowman PF, Sallan L, Friedman M, Kaschner K, Garilao C, Near TJ, Coll M, Alfaro ME. 2018. An inverse latitudinal gradient in speciation rate for marine fishes. Nature 559:392–395.

Reynalte-Tataje DA, Lopes CA, de Ávila-Simas S, Garcia JRE, Zaniboni-Filho E. 2013. Artificial reproduction of neotropical fish: Extrusion or natural spawning? Natural Science 5:1–6.

Rice AN, Soldevilla MS, Quinlan JA. 2017. Nocturnal patterns in fish chorusing off the coasts of Georgia and eastern Florida. Bulletin of Marine Science 93:455–474.

Riera, A., R. A. Rountree, L. Agagnier, F. Juanes. 2020. Sablefish (*Anoplopoma fimbria*) produce high frequency rasp sounds with frequency modulation. Journal of the Acoustical Society of America 147:2295–2301.

Rigley L, Marshall JA. 1971. Sound production by the loach *Botia berdmorei* (Pisces, Cobitidae). American Zoologist 11:632.

Rountree RA, Juanes F. 2010. First attempt to use a remotely operated vehicle to observe soniferous fish behavior in the Gulf of Maine, Western Atlantic Ocean. Current Zoology 56:90–99.

Salmon M, Winn HE. 1966. Sound production by priacanthid fishes. Copeia 1966:869–872.

Salmon M, Winn HE, Sorgente N. 1968. Sound production and associated behavior in triggerfishes. Pacific Science 22:11–20.

Schachner G, Schaller F. 1981. Schallerzeugung und Schallreaktionen beim Antennenwels (Mandim) *Rhamdia sebae sebae* Val. Zoologische Beitraege 27:375–392.

Scholz K, Ladich F. 2006. Sound production, hearing and possible interception under ambient noise conditions in the topmouth minnow *Pseudorasbora parva*. Journal of Fish Biology 69:892–906.

Shimizu T. 1977. Comparative morphology of the expanded epipleural and its associated structures in four species of the Trachichthyidae. Japanese Journal of Ichthyology 23:192–198.

Širović A, Demer DA. 2009. Sounds of captive rockfishes. Copeia 2009:502–509.

Skelton PH, Skelton PH. 1993. A Complete Guide to the Freshwater Fishes of Southern Africa. Halfway House, South Afrida: Southern Book Publishers.

Sorensen W. 1890. Om Forbeninger i Svømmeblæren, Pleura og Aortas Virg og Sammensmeltning deraf med Hvirvelsøjlen særlig hos Siluroiderne, samt de saakaldte Weberske Knoglers Morphologi. Det Kongelige Danske Videnskabernes Selskabs Skrifter 6 Række: Naturvidenskabelig og Mathematisk Afdeling 6:65–152.

Sörensen W. 1884. Om Lydorganer hos Fiske: En physiologisk og comparativ-anatomisk Undersøgelse. Copenhagen: V. Thaning & Appels.

Sörensen W. 1894-1895. Are the extrinsic muscles of the air-bladder in some Siluroidae and the “elastic spring” apparatus of others subordinate to the voluntary production of sounds? What is, according to our present knowledge, the function of the weberian ossicles? A contribution to the biology of fishes. Journal of Anatomy and Physiology 29:109–139; 205-229; 399-423; 518-552.

Spanier E. 1979. Aspects of species recognition by sound in four species of damselfishes, genus *Eupomacentrus* (Pisces: Pomacentridae). Zeitschrift für Tierpsychologie 51:301–316.

Straight CA, Freeman BJ, Freeman MC. 2014. Passive acoustic monitoring to detect spawning in large-bodied catostomids. Transactions of the American Fisheries Society 143:595–605.

Takayama M, Onuki A, Yosino T, Yoshimoto M, Ito H, Kohbara J, Somiya H. 2003. Sound characteristics and the sound producing system in silver sweeper, *Pempheris schwenkii* (Perciformes: Pempheridae). Journal of the Marine Biological Association of the UK 83:1317–1320.

Tavolga WN. 1958. Underwater sounds produced by males of the blenniid fish, *Chasmodes bosquianus*. Ecology 39:759–760.

Tavolga WN. 1964. Sonic characteristics and mechanisms in marine fishes. In: Tavolga WN, editor. Marine Bio-acoustics. New York: Pergamon Press. p. 195–211.

Taylor M, Mansueti RJ. 1960. Sounds produced by very young crevalle jack, Caranx hippos, from the Maryland seaside. Chesapeake Science 1:115–116.

Tower RW. 1908. The production of sound in the drumfishes, the sea-robin and the toadfish. Annals of the New York Academy of Sciences 18:149–180.

Tricas TC, Boyle KS. 2014. Acoustic behaviors in Hawaiian coral reef fish communities. Marine Ecology Progress Series 511:1–16.

Tricas TC, Boyle KS. 2015. Diversity and evolution of sound production in the social behavior of *Chaetodon* butterflyfishes. Journal of Experimental Biology 218:1572–1584.

Uchida K. 1934. On the sound-producing fishes of Japan. Nihon Gakujitsu Kyokai Hokoku 9:369–375.

Walls PD. 1964. The anatomy of the sound producing apparatus of some Australian teleosts Honors Thesis. Brunswick, ME: Bowdoin College.

Walters V. 1960. The swimbladder of *Velifer hypselopterus*. Copeia 1960:144–145.

Weitzman SH. 1954. The osteology and the relationships of the South American Characid fishes of the sub-family Gasteropelecinae. Stanford Ichthyological Bulletin 4:213–263.

Wilson B, Batty RS, Dill LM. 2004. Pacific and Atlantic herring produce burst pulse sounds. Biology Letters 271:S95–S97.

Winn HE, Stout JF. 1960. Sound production by the satinfin shiner, *Notropis analostanus*, and related fishes. Science 132:222–223.

Yabe M. 1985. Comparative osteology and myology of the superfamily Cottoidea (Pisces, Scorpaeniformes) and its phylogenetic classficiation. Memoirs of the Faculty of Fisheries Hokkaido University 32:1–130.

Yabe M, Uyeno T. 1996. Anatomical description of *Normanichthys crockeri* (Scorpaeniformes, Incertae sedis: Family Normanichthyidae). Bulletin of Marine Science 58:494–510.

Yamato K, Matsuo I, Takahashi R, Matsubara N, Yasuma H. 2018. Call localization of fat greenling *Hexagrammos otakii* using two stereo underwater recorders. Marine Technology Society Journal 52:129–138.

Yokoyama K, Kamei Y, Toda M, Hirano K, Iwatsuki Y. 1994. Reproductive behavior, eggs, and larvae of a caesionine fish, Pterocaesio digramma, observed in an aquarium. Japanese Journal of Ichthyology 41:261–274.

